# Non-apoptotic caspase events and Atf3 expression underlie direct neuronal differentiation of adult neural stem cells

**DOI:** 10.1101/2024.03.09.584233

**Authors:** Frédéric Rosa, Nicolas Dray, Laure Bally-Cuif

## Abstract

Neural stem cells (NSCs) are key physiological components of adult vertebrate brains, generating neurons over a lifetime. In the adult zebrafish pallium, NSCs persist at long term through balanced fate decisions that include direct neuronal conversions, i.e., delamination and neurogenesis without a division. The characteristics and mechanisms of these events remain unknown. Here we reanalyze intravital imaging data of adult pallial NSCs and observe shared delamination dynamics between NSCs and committed neuronal progenitors. In a candidate approach for mechanisms predicting NSC decisions, we build an NSC-specific genetic tracer of Caspase3/7 activation (Cas3*/Cas7*) *in vivo* and show that non-apoptotic Cas3*/7* events occur in adult NSCs and are biased towards neuronal conversion under physiological conditions. We further identify the transcription factor Atf3 as necessary to express this fate. Finally, we show that the Cas3*/7*/Atf3 pathways are part of the processes engaged when NSCs are recruited for neuronal regeneration. These results provide evidence for the non-apoptotic caspase events occurring in vertebrate adult NSCs and link these events with the NSC fate decision of direct conversion, important for long-term NSC population homeostasis.

## Introduction

Stem cell (SC) fate decisions orchestrate adult organ maintenance and SC population renewal over a lifetime. These decisions often occur in link with SC division; as a result, much work has been devoted to understanding SC proliferation frequency and daughter cell fate control. However, it has also been reported in a few instances, notably in the brain and muscle, that a differentiated fate could be directly acquired from an adult SC in the absence of a division event under physiological conditions (Barbosa et al., 2015; Bjornson et al., 2012; Dray et al., 2015; Ismaeel et al., 2023; Mourikis et al., 2012). These events described as “direct conversion” have remained understudied. Their characteristics are not precisely described, and it remains largely unknown how they are controlled and what their physiological relevance may be.

In the adult vertebrate brain, neural stem cells (NSCs) are radial astroglia, which are mostly quiescent and divide at low frequency (Chaker et al., 2016; Lampada and Taylor, 2023; Obernier and Alvarez-Buylla, 2019; Urbán et al., 2019; Yeh et al., 2023). In mouse, NSCs are specialized SC that can generate differentiated neurons and astrocytes, both of which are molecularly distinct from NSCs (Beckervordersandforth et al., 2010; Dulken et al., 2017; Zywitza et al., 2018). In zebrafish, NSCs co-express markers of mature astrocytes (e.g., Glial fibrillary acidic protein -Gfap-, or Glutamine synthase -GS-) and progenitors (e.g., *her4* genes) during their quiescence phase (Cosacak et al., 2019; Morizet et al., 2024). Upon division, they generate NSCs and/or committed neuronal progenitors (NPs) (negative for Gfap and/or GS) that differentiate into neurons (Dirian et al., 2014; Furlan et al., 2017; Kroehne et al., 2011; Mancini et al., 2023). In both models, genetic clonal tracing and/or intravital imaging also provided evidence for individual NSCs acquiring a neuronal fate while no cell division was detected. For example, in the mouse dentate gyrus, around 10% of clones issued from *Nestin:Cre*-mediated tracing of individual NSCs consist of single neurons after one month (Bonaguidi et al., 2011), suggesting a direct conversion -although the interpretation of these results can be confounded by cell death-. In the zebrafish pallium, intravital imaging reveals direct neuronal differentiation occurring from *her4.3-* positive or *gfap*-positive NSCs that do not express proliferation markers and/or have not divided since at least 14 days (4 imaging time points, 14-16 days) (Barbosa et al., 2015; Dray et al., 2015; Than-Trong et al., 2020). Longitudinal imaging in these cases avoids ambiguity as no cell death events were observed, but the occurrence of direct conversion events remains difficult to quantify at population scale as the duration of imaging is short relative to NSC division frequency. *her4:ERT2CreERT2*-mediated clonal tracing and modeling, performed on a dataset spanning over >530 days, predict that direct conversions represent 25% of fate decisions (Than-Trong et al., 2020), and are crucial to maintain homeostasis of the NSC population. Indeed, a long-term equilibrium of NSC numbers is observed *in situ* and results from balancing NSC gains (through amplifying divisions) and losses (through symmetric neurogenic divisions and direct differentiation events) (Than-Trong et al., 2020).

In the line of our current work aiming to identify predictors and mechanisms of NSC fate decisions in vivo, we focused on NSC direct conversions and challenged whether they could be connected with non-apoptotic caspase events. Caspases are site-specific proteases of the programmed cell death system. In addition to the execution of cell death, however, caspases are increasingly recognized to participate in non-apoptotic events during development and homeostasis in a variety of tissues and organisms (reviewed in (Abdul-Ghani and Megeney, 2008; Burgon and Megeney, 2018). In particular, there is a frequent association of non-apoptotic caspase events with cell fate decisions, including in some SCs e.g. ES cells or hematopoietic SCs (Fujita et al., 2008; Janzen et al., 2008). In the nervous system, non-apoptotic caspase recruitments modulate dendritic pruning, developmental circuit maturation and axonal pathfinding, to cite a few (Unsain and Barker, 2015). In embryonic neurospheres in vitro, blocking these events limits neuronal differentiation and neurite extension (Fernando et al., 2005). Finally, in the embryonic peripheral nervous system in Drosophila, non-apoptotic caspase-mediated cleavage of signaling pathway effectors limit neural progenitor proliferation to control neuronal production (Colon-Plaza and Su, 2022; Kuranaga et al., 2006). Non-apoptotic caspase events involve activation of the direct caspase pathway converging onto the activation of effector Caspases 3 and 7 (activated forms noted as Cas3*, Cas7*) by proteolytic cleavage, but can be tracked using Cas3*/Cas7* sensors. For example, in Drosophila, CasExpress and CaspaseTracker liberate Gal4 upon cleavage at the Cas3*/Cas7* canonical site DEVD, to drive lineage labeling when caspase activation is not followed by cell death (Ding et al., 2016; Tang et al., 2015). These studies revealed multiple cells surviving an early caspase event under physiological conditions. Non-apoptotic Cas3* or Cas7* can be seen as true death reversals (referred to as anastasis) (Sun et al., 2017; Tang and Tang, 2018). Rather than interrupted apoptosis, they may also signify the use of common pathways to trigger cell remodeling shared between apoptosis and cellular decisions. Whether non-apoptotic caspase events take place in adult NSCs is unknown.

In this paper, we proceed in two initially independent steps: first, we attempt to characterize the morphodynamic features of direct conversion events; second, we address whether non-apoptotic caspase events take place in NSCs in the adult vertebrate brain, and if so, whether these events encode specific fate decisions. We use as model the zebrafish adult pallium, the cartesian construction of which allows inferring the birthdate and origin of any neuron from its position at the time of analysis (Furlan et al., 2017). We generate an inducible, selective and stable Cre-mediated Cas3*/Cas7* sensor, *Cas^CRE^Atlas*, which allows long-term fate tracing of non-apoptotic Cas3*/Cas7* events occurring in NSCs. We find that such events do take place in NSCs and, further, that they are biased towards fate choices of direct neuronal generation under physiological conditions. Combining gain- and loss-of-function experiments, we identify the stress-induced transcription factor Atf3 as necessary for this NSC fate in vivo. Finally, we show that Cas3*/Cas7* events are also induced in response to lesion. Together, these results highlight the occurrence of non-apoptotic caspase events in NSCs during pallial development and adult life, and demonstrate their relevance for a specific NSC fate decision involved in the physiological homeostasis of adult NSC populations.

## Results

### Direct conversion events and post-division delaminations share morphodynamic features

Using intravital imaging of the whole NSC population in the adult zebrafish pallium, NSC direct conversion events were previously defined as the loss of expression of the NSC marker Tg(*gfap:dTomato*) accompanied with delamination from the pallial ventricular layer in the absence of visible division during the preceding 4 imaging time points (14 to 16 days) (Than-Trong et al., 2020). The choice of this duration was based on the observation that, when a neurogenic division is visible in a movie, 85 +/- 10% of neuron-fated daughter cells visibly express their fate (loss of *gfap*:dTomato) within the 10-12 days post-division (Dray et al., 2021; Than-Trong et al., 2020). Such events are followed by expression of the neuronal differentiation marker HuC/D and the presence of a neuronal process (Barbosa et al., 2015).

To characterize the morphodynamic features of these events and determine whether and how they contrast with delamination and differentiation events occurring as a result of neurogenic divisions (hereafter referred to as “post-division delaminations”), we exploited the intravital imaging movies acquired in four 3-month post-fertilization (mpf) adults in the *Casper*;Tg(*gfap:ZO1- mKate2*);Tg(*deltaA:egfp*) background (Mancini et al., 2023). This dataset contains a total of 828 NSCs filmed over 39 to 43 days every 2 to 3 days in 4 pallial hemispheres (from 4 different fish) and reveals NSC apical surfaces (ZO1-mKate2) and a neurogenic fate (*deltaA* expression). We focused here on delaminations, which were not previously analyzed. We defined as “delamination termination” the first time point with no identifiable ZO1-mKate2-negative apical surface (within the resolution of our intravital two-photon imaging), i.e., the moment of apical closure (Fig.1A); this time point is then followed by the reestablishment of apicobasal boundaries between remaining cells. When no division is visible along the track, we refer to “time point 0” (tp0) as the first imaging time point of the movie (Fig.1A, top); in that case, we excluded delaminations occurring less than 4 time points (14-16 days) from the start of each movie, to avoid missing recent division events. When a division visible along the track, tp0 is the first time point post-division (Fig.1A, bottom). With these criteria, the dataset includes 83 delamination events. 54 of these delaminations terminated after at least 4 imaging time points (14-16 days) without visible division, and 29 followed a visible division during the previous 3 imaging time points or less (<= 13 days). In the first category, the time from tp0 to delamination termination varies between 14 and 41 days. Two examples are illustrated in Fig.1B, displaying delamination events that occur without detectable division during the previous 23 days (example 1) and 35 days (example 2).

**Figure 1.**
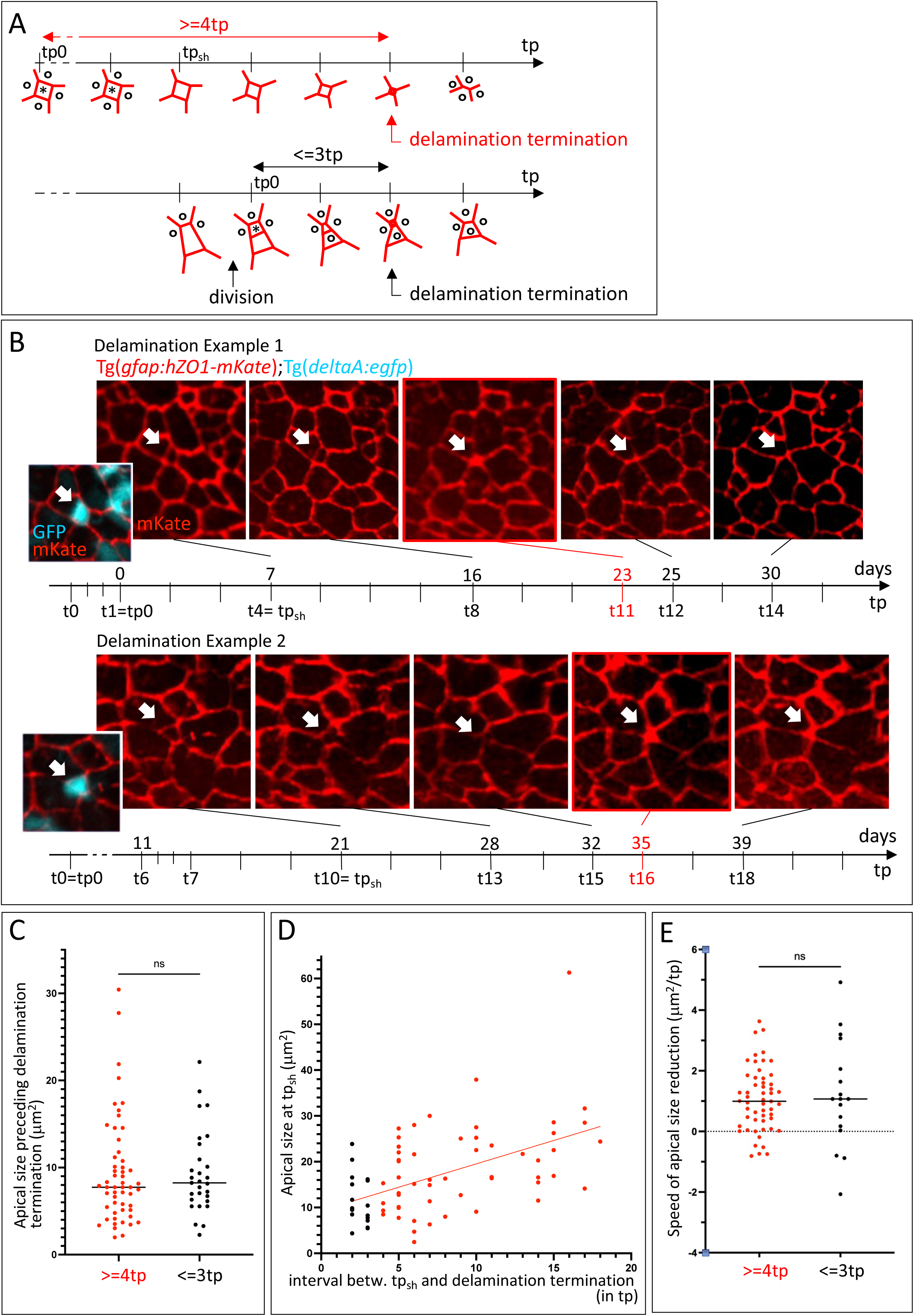
Intravital imaging permits the morphodynamic characterization of delamination events in the adult pallium. **A.** Schematics of a delamination events observed using intravital imaging in Tg(*gfap:ZO1-mKate2*) 3mpf adults. Horizontal arrows: imaging time points (tp). In each series, the apical surface of a delaminating cell (*) is depicted, with 4 non-delaminating neighbors (°) (apical view, red segments to ZO1-positive cellular interfaces). Delamination is considered terminated when no ZO1- negative apical domain is visible. **Top**: the tracking encompasses at least 4 imaging time points (tp) (14- 16 days) without visible division prior to delamination termination. In that case, tp0 is the first time point of the movie. Tracks with no division but delamination terminating less than 4 time points after the start of imaging are not considered. tp_sh_ is the time point at which apical size shrinkage is initiated. **Bottom**: a division event takes place 3 tp or less (13 days or less) before delamination termination. In that case, tp0 if the first time point post-division. **B. Top**: Snapshot series of two delamination events recorded from a *Casper*;Tg(*gfap:hZO1-mKate*);Tg(*deltaA:egfp*) 3mpf adult (fish named Outi, raw data in (Mancini et al., 2023)). Only the ZO1-mKate channel is shown (red). White arrows to delaminating cells, the first tp illustrated is the onset of apical area shrinkage (tp_sh_), the delamination termination tp is framed in red, the vertical time arrows indicate imaging tp (left) and corresponding days (right). The apical area of cells in examples 1 and 2 are 20.1 μm2 and 22.9 μm2, respectively. **Bottom left insets**: Images including the *deltaA*:eGFP channel (cyan) at the onset of shrinkage. **C-E.** Quantified dynamic parameters of delaminations occurring after at least 4 time points (14-16 days) without division (red) and delaminations occurring 3 time points or less (<= 13 days) post-division (black). **C.** Apical surface area at the tp preceding delamination termination. Mann Whitney: test, p-value: 0.661. **D.** Apical surface area at tp_sh_ as a function of the time elapsed between tp_sh_ and delamination termination. Linear regression, R squared = 0.256. **E.** Speed of apical size reduction (shrinkage speed), calculated from the tp_sh_ until delamination termination. Mann Whitney: test, p-value: 0.849.

We next analyzed the measurable morphometric parameters of delaminations, comparing those occurring after at least 4 time points without division vs 3 time points or less post-division (red and black on Fig.1C-E, respectively). Cells in these two categories did not differ significantly in their apical area at the time point preceding delamination termination (on average, 9.4 μm^2^, +/- 6.1 μm^2^ SD and 9.3 μm^2^, +/- 4.8 μm^2^ SD respectively) (Fig.1C), nor in their expression of *deltaA* (100% *deltaA^pos^* cells in both cases) (examples in Fig.1B, bottom). There was a moderate correlation between apical surface area at the onset of apical surface shrinkage (tp_sh_) and the duration of apical shrinkage during the delamination process (cells starting with a larger apical surface area taking longer to delaminate) (Fig.1D). When normalized over the duration of shrinkage until delamination termination, shrinkage rates appeared similar between the two categories (∼1μm^2^ per time point) (Fig.1E). Together, these results characterize the quantitative and molecular features of the delamination process during adult pallial neurogenesis. They reveal its comparable dynamics irrespective of time post-division or apical surface area at the start.

The dataset does not allow to directly recognize direct conversions, as NPs can be misled for NSCs. Indeed, ZO1-mKate2 expressed by NSCs also surrounds intermingled NPs and makes them tractable. NSCs have on average larger apical surface areas than NPs (with medians at 100μm^2^ vs 20μm^2^, respectively), but the distributions are broad and overlap ((Mancini et al., 2023) and N. Dray, unpub.). To estimate the percentage of NSCs in our dataset, we considered the two possible NP configurations: (i) isolated NPs surrounded by NSCs, and (ii) NP clusters (e.g., an NP doublet generated by the symmetric neurogenic division of an NSC or an NP, which will appear as a single apical surface surrounded by ZO1-mKate) (Fig.S1A). To quantify NP clusters, we counted the number of ZO1-mKate surfaces tracked in intravital imaging that in fact contained adjacent NPs, by comparing our live data with immunohistochemistry for ZO1 on the same specimen fixed at the end of the film (Fig.S1A). The percentage of surrounded NP clusters increased as the ZO1-mKate surface area decreased (Fig.S1B), as logically expected from the fact that NPs have smaller apical surface areas than NSCs. With an apical surface area of 30μm^2^ or less at tp_sh_ (which is a large majority of recorded events, 80 over 83 events, 96%) (see below and Fig.1D), 21.4% of ZO1-mKate surfaces were NP clusters (Fig.S1B). To count isolated NPs, an NSC marker was necessary and we used fixed Tg(*gfap:dTomato*) pallia immunostained for Tomato and ZO1. With an apical surface area of 30μm^2^ or less, 29.3% of surfaces were isolated NPs (Fig.S1B). Together, we conclude that approximately half of the ZO1-mKate surfaces of 30μm^2^ or less recorded in intravital imaging are NSCs (Fig.S1C). This makes it likely that the morphometric features measured above (Fig.1) also apply to NSC delaminations (referred to as direct conversions).

### Non-apoptotic Cas3*/Cas7*-mediated cleavage events take place during pallium development and homeostasis

In our search for predictive parameters of NSC decisions, including direct conversions, we next considered non-apoptotic Caspase events. To determine whether non-apoptotic proteolytic events at the DEVD Cas3*/Cas7* target site (Chéreau et al., 2003; Talanian et al., 1997) occur in the developing and adult zebrafish pallium, we designed a heritable lineage tracer of DEVD cleavage in NSCs. The driver transgenic line, Tg*(her4:mCD8-DEVD-V5-Cre)*, produces a Cre recombinase (fused to a V5 tag) tethered to the plasma membrane via a DEVD site, expressed under control of the NSC *her4.3* regulatory elements (Yeo et al., 2007) (Fig.2A). When Tg*(her4:mCD8-DEVD-V5-Cre)* fish are crossed into the Tg*(μact:lox-stop-lox-hmg2bmCherry)* reporter background (Wang et al., 2011) (this double background is hereafter referred to as *Cas^CRE^Atlas*), non-apoptotic cleavage events at the DEVD site in *her4*-expressing cells are expected to trigger Cre-mediated recombination and the permanent expression of Hmg2bmCherry in all progeny cells. A Tg(*her4:mCD8-GSGC-V5-Cre*) driver, immune to Cas3*/Cas7*, was used as control.

**Figure 2.**
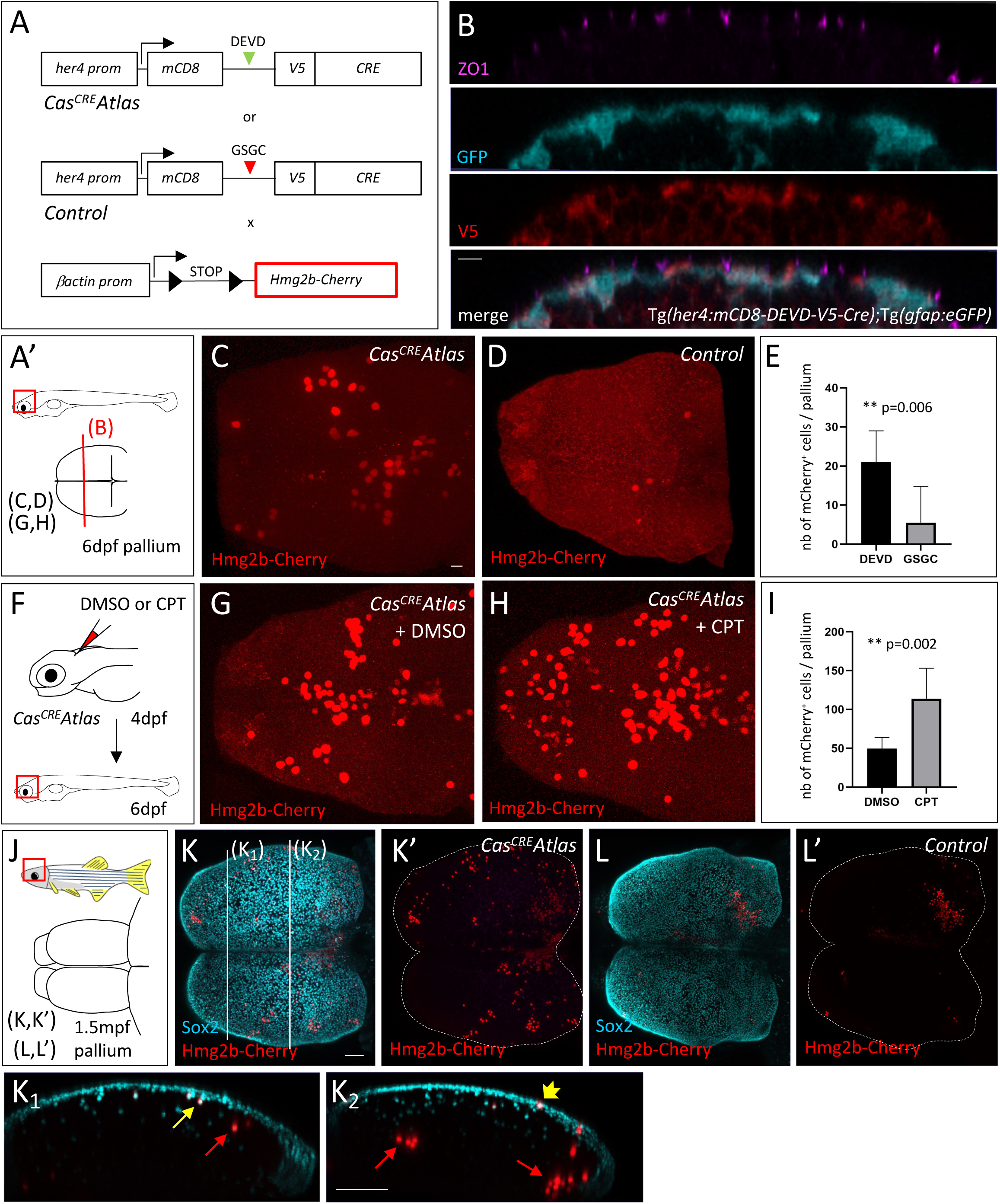
The *Cas^CRE^Atlas* approach reveals non-apoptotic caspase-triggered events during pallium development and homeostasis. **A.** The *Cas^CRE^Atlas* principle combines a NP/NSC-specific driver line (top and middle) expressing the Cre recombinase upon Cas3*/Cas7* cleavage with a reporter line (bottom) expressing *hmg2bmCherry* upon Cre recombination. mCD8: membrane anchor; V5: antigen segment (tag); DEVD: canonical Cas3*/Cas7* cleavage site; GSGC: mutated non-productive cleavage site; *her4* prom: *her4* regulatory elements. The *Tg(her4:mCD8-DEVD-V5-Cre);Tg(-act;lox-stop-lox- hmg2bmCherry)* background is referred to as *Cas^CRE^Atlas*, and *Tg(her4:mCD8-GSGC-V5-Cre);Tg(bact;lox-stop-lox-hmg2bmCherry)* as *Control*. All analyses are conducted in double heterozygotes. **A’.** Schematic of analyses in cross-sectioned or whole-mount pallia at 6dpf. **B.** Expression of the *Cas^CRE^* driver is confined to *gfap:eGFP*-positive cells. Cross sections through the pallium of *Tg(her4:mCD8-DEVD-V5- Cre);Tg(gfap;eGFP)* double transgenic larvae at 6dpf with triple immunostaining for GFP (NP/NSC), ZO1 (tight junctions at the apical surface of pallial ventricular cells) and V5 (transgene tag) (color coded), showing expression of V5 (membrane-anchored) and GFP in the same cells. **C-E.** Hmg2bmCherry whole-mount immunohistochemistry on *Cas^CRE^Atlas* (C) and *Control* (D) pallia at 6dpf (dorsal views, z projections, anterior left), and quantification of positive cells (E) (*Cas^CRE^Atlas*: n=4 brains, *Control*: n=10 brains, Mann-Whitney test). **F-I.** *Cas^CRE^Atlas* activity is induced by transient exposure to Camptothecin (CPT), an apoptosis inducer. **F.** Schematic of experiment: CPT (20μM) or DMSO are injected through the hindbrain into the neural tube ventricle in *Cas^CRE^Atlas* larvae at 4dpf, and Hmg2bmCherry is analyzed at 6dpf. **G-I.** Hmg2bmCherry whole-mount immunohistochemistry at 6dpf (dorsal views, z projections, anterior left) (G, DMSO; H, CPT) and quantification (I) (DMSO and CPT: n=6 brains each, Mann-Whitney test). **J-L’.** Hmg2bmCherry expression in *Cas^CRE^Atlas* (K) and *Control* (L) pallia at 1.5mpf. **J.** Schematic of analyses in whole-mount pallia at 1.5mpf. **K-L’.** Double whole-mount immunohistochemistry for Sox2 (NPs, NSCs and some freshly born neurons) and Hmg2bmCherry (K,K’,L,L’: dorsal views, z projections, anterior left, channels color-coded; K_1_,K_2_: cross sections at the levels indicated in K, dorsal up). Red arrows: Sox2-negative neurons, yellow arrow: Sox2-positive neurons, short yellow arrow: Sox2-positive NSC. Scale bars: B,C, D, G, H: 10µm, K-L’, K1-2: 50µm.

To validate the approach, we first analyzed the expression of the *her4:mCD8-DEVD-V5-Cre* transgene and the inducibility and selectivity of *Cas^CRE^Atlas* tracing in the larval pallium at 6 days post-fertilization (dpf). Immunohistochemistry in the double transgenic Tg*(her4:mCD8-DEVD-V5-Cre)*;Tg(*gfap:eGFP*) background showed that V5-Cre expression is faithfully restricted to NPs in the larval pallium (Fig.2B). Next, we used whole-mount immunohistochemistry to test whether Hmg2bmCherry-positive cells were present in the pallium of 6dpf *Cas^CRE^Atlas* versus control larvae. While Hmg2bmCherry events were rare in control pallia -possibly reflecting a low affinity of Cas3*/Cas7* for non-canonical sites, or cleavage of GSGC by another protease at low frequency-, Hmg2bmCherry-positive cells were easily detectable and significantly more numerous in *Cas^CRE^Atlas* pallia (Fig.2A’,2C-E). Finally, we tested whether *Cas^CRE^Atlas* was responsive to activation of the caspase cascade. 4dpf larvae were transiently subjected to the apoptosis inducer Camptothecin (CPT) (Ikegami et al., 1999) injected into the hindbrain ventricle, then chased until 6dpf. CPT significantly increased the number of Hmg2bmCherry-positive cells in *Cas^CRE^Atlas* larvae compared to control DMSO injections (Fig.2F-I). These results together demonstrate the selective and sensitive tracing of non-apoptotic Caspase events by *Cas^CRE^Atlas* in the zebrafish pallium *in vivo*, and for the occurrence of such events under physiological conditions at least at larval stages.

The zebrafish pallium follows an outside-in neuronal generation pattern, whereby neurons generated at early stages end up in deep parenchymal locations whereas late-born neurons are positioned more superficially, close to the active NSC monolayer (Furlan et al., 2017). Tangential neuronal migration does not take place, and these properties together allow using the position of a neuron as a spatio- temporal signature to infer its generation time and approximative ventricular region of origin. We used these properties to characterize and interpret the pattern of Hmg2bmCherry cells at juvenile and adult stages in *Cas^CRE^Atlas* animals (Fig.2J and see below). At these stages, neurogenesis follows the sequence NSCs (*her4*^pos^ or *gfap*^pos^; Sox2^pos^) > NPs (*her4*^neg^ and *gfap*^neg^; Sox2^pos^) > neurons (Than-Trong et al., 2020). At 1.5mpf, a few deep groups of Hmg2bmCherry cells could be occasionally seen in control pallia (Fig.2L,L’), resulting from early non-specific activation events as described at 6dpf. In contrast, the Hmg2bmCherry pattern of *Cas^CRE^Atlas* 1.5mpf pallia was very different, with numerous small groups of labeled cells, some located in superficial pallial layers (Fig.2K,K’). Immunohistochemistry for Sox2 - which also labels some recently born neurons- was used as a landmark and revealed deeply located (old) Sox2-positive neurons as well as more superficial (recent) events, with staining of either freshly born neurons, NSCs or NPs (Fig.2K_1_,K_2_). We interpret this pattern as reflecting Cre recombination events that occurred at different time points (from older to more recent) in *her4*-expressing cells.

### The Cas3*/Cas7*-driven lineage is biased towards direct neuronal differentiation

Cas3/Cas7 activation in *her4*-positive cells may reflect transient events unrelated to NSC decisions or may correlate with specific behaviors. To address this, we compared NSC fate in *Cas^CRE^Atlas* labeled clones with *her4*-mediated NSC fate tracing in the juvenile to young adult pallium (Fig.3A). Because neurons generated from pallial NSCs stack in age-related layers (Furlan et al., 2017), we established a temporal landmark across pallial depth, using a BrdU pulse at 1mpf, to label neurons born at that stage. This allowed to select *Cas^CRE^Atlas* events having occurred between 1mpf and the stage of analysis (2mpf): these events generated Hmg2bmCherry-positive cells located above the BrdU landmark (Fig.3B). In parallel, clonal recombination events were induced in *Tg(her4:ERT2CreERT2);Tg(μact:lox- stop-lox-eGFP)* double transgenic animals just prior to the BrdU pulse, and analyzed at 2mpf as well.

**Figure 3.**
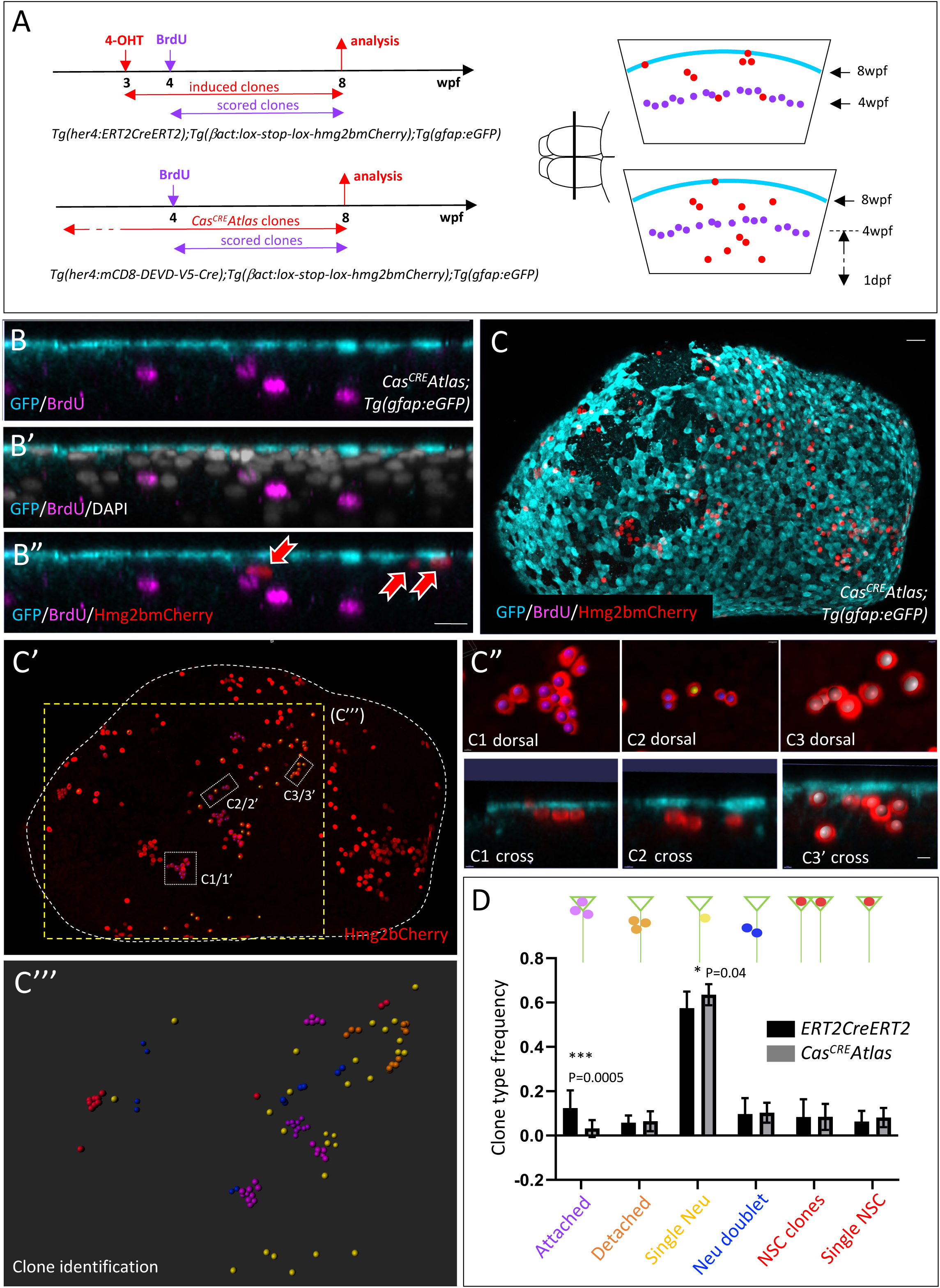
*Cas^CRE^Atlas*-driven clones in pallial NSCs display a biased fate. **A.** Schematic of the experiment. *Tg(her4:ERT2CreERT2);Tg(μact:lox-stop-lox-hmg2bmCherry);Tg(gfap:eGFP)* (top) and *Cas^CRE^Atlas;Tg(gfap:eGFP)* (bottom) triple transgenic fish are used to compare NSC fate between unbiased and Cas3*-driven clones, respectively. Left panels: temporal sequence of 4-OHT induction, BrdU pulse labeling, and analysis. Right panels: schematic pallial cross sections showing the relative positions of the NSC layer (cyan), BrdU-labeled neurons (magenta) and Hmg2bmCherry-positive clones (red) in each genotype. We focused on *Cas^CRE^Atlas* clones generated post-4 weeks-post-fertilization (wpf), i.e. situated between the layer of BrdU-positive neurons and the pallial ventricle. **B.** Cross section in a *Cas^CRE^Atlas;Tg(gfap:eGFP)* triple transgenic fish, processed for immunohistochemistry to reveal GFP (cyan, NSCs), BrdU (magenta) and Hmg2bmCherry (red) exemplifying the cartoon in A, bottom. Red arrows to Hmg2bmCherry-positive cells. **C-C’’’.** Whole-mount *Cas^CRE^Atlas;Tg(gfap:eGFP)* 2mpf pallium stained for GFP, BrdU and Hmg2bmCherry (dorsal view, anterior left). C,C’: channels color-coded; C”: examples of several clones (C1-C3, indicated on C’), with color-coded segmentation, views from dorsal (C1-C3) and in optical cross sections (C1’-C3’) -not all cells are visible in the latter case-. Clones in C1 are “attached” (magenta), clones in C2 are “single Neu” (1 clone, yellow) and “Neu doublets” (2 clones, blue), clones in C3 are “single Neu” (2 clones, yellow) and “detached” (1 clone, orange); C’’’: Imaris segmentation of unambiguously identifiable clones in the area shown in C’ (yellow dotted square). **D.** Quantification of the frequency of the different Hmg2bmCherry-positive clone types (represented schematically along the z axis; green triangles: NSCs; colored dots: Hmg2bmCherry-positive cells) generated from *her4*-positive NSCs between 4 and 8 wpf in the *Tg(her4:ERT2CreERT2);Tg(μact:lox-stop- lox-hmg2bmCherry);Tg(gfap:eGFP)* and *Cas^CRE^Atlas;Tg(gfap:eGFP)* backgrounds (black and gray bars, respectively) (n=5 brains for each genotype, 344 clones for unbiased clonal tracing (4-OHT treatment), n=360 clones for *Cas^CRE^Atlas* tracing). Statistical analysis performed by contingency Chi-square test, 95% confidence. Scale bars: B,B’: 15µm, C, C’, C”: 30 µm, C3-8: 10µm.

We considered that Hmg2bmCherry-positive cells belong to the same clones when they were separated by less than a 2-cell diameter distance. This is based on previous observations documenting the absence of cell migration or death of *her4* progeny cells in the zebrafish pallium (Furlan et al., 2017; Webb et al., 2009). The composition of clones was assessed in 3D in whole-mount pallia using immunohistochemistry for *gfap*:eGFP (NSCs). GFP-negative cells include NPs and neurons and, for simplicity, were labeled as neurons (Fig.3C-C”). As expected, we found that unbiased clonal labeling generates a variety of clone types (Fig.3D). Specifically, single NSCs are interpreted as NSCs that remained quiescent since the labeling pulse, NSC doublets as resulting from an amplifying NSC/NSC division (red in Fig.3D), neuron doublets from -most likely- a symmetric neurogenic division, and single neurons from a direct differentiation event (blue and yellow, respectively, in Fig.3D). The lineage tree of clones composed of 3 cells or more cannot be resolved, and these clones were simply classified as “attached” or “detached” depending on their link or not with the NSC layer (magenta and orange, respectively, in Fig.3D). They reflect ongoing, or terminated, productive neurogenic events, respectively. These fates together are in qualitative and quantitative agreement with the different division modes and fates reported in previous work (Than-Trong et al., 2020). We found Cas3*Cas7* events associated with most clone types; however, attached clones were virtually absent, while the proportion of single neurons was increased (Fig.3D). Thus, non-apoptotic Cas3*/Cas7* events in pallial NSCs correlate with a bias of NSC fate towards direct neuronal differentiation. Finally, this analysis allowed to estimate that, on average, 50 *Cas^CRE^Atlas* events occur during 4 weeks within an hemipallium (i.e., an area of approximately 2000-2500 *her4*^pos^ NSCs).

### The Atf3 transcription factor, a mediator of anastasis, is expressed in a subset of scattered delaminating cells at the adult pallial ventricle at any time

We next aimed to identify molecular events involved in promoting the direct differentiation fate of pallial NSCs. The results above are correlative, but suggestive. Hence, we used a candidate approach searching among known mediators of the non-apoptotic caspase event known as anastasis (Sun et al., 2017; Tang et al., 2017). A common molecular signature of anastasis was identified in various mammalian cell types (Sun et al., 2017; Tang et al., 2017). The orthologs of many genes up-regulated at an early stage of anastasis fate reversal appeared expressed in a subset of pallial NSCs under physiological conditions, as revealed in our scRNAseq dataset (Morizet et al., 2024). Among these genes (Fig.S2A), we chose to focus on the transcription factor-encoding gene *atf3*, as one of the top genes induced in mammalian cells, and because its scRNAseq expression was detectable in a low number of pallial quiescent NSCs in a manner unlinked with clustering (Fig.4A). A low cell number is in agreement with the low frequency of NSC fate decisions at any given time, considering the slow dynamics of the NSC population (Dray et al., 2021; Than-Trong et al., 2020). Sparse expression was confirmed using whole-mount in situ hybridization (ISH), which revealed scattered *atf3*-positive cells across the pallial surface (Fig.4B). Some areas exhibited a higher concentration of *atf3*-positive cells (such as the posterior and dorsomedial pallium), but at small scale the *atf3* pattern is not exactly identical in each of these areas in the two hemispheres. Finally, we used the *Tg(gfap:eGFP)* background and fluorescent chromogenic ISH to address the morphology of *atf3*-positive cells. Cross-sections, or horizontal optical focus at progressively deeper z planes from the ventricular surface, showed that *atf3*-positive;GFP-positive cells have a delaminating profile, with their nuclei often located in deeper positions than the majority of NSCs (Fig.4C,D). Two *atf3* transcripts were recovered in the adult pallium, corresponding to alternative splicing events predicted to encode long and short protein isoforms that differ in their N-terminus (Fig.4E). Our ISH used the *atf3-L* probe and does not distinguish between these isoforms.

**Figure 4.**
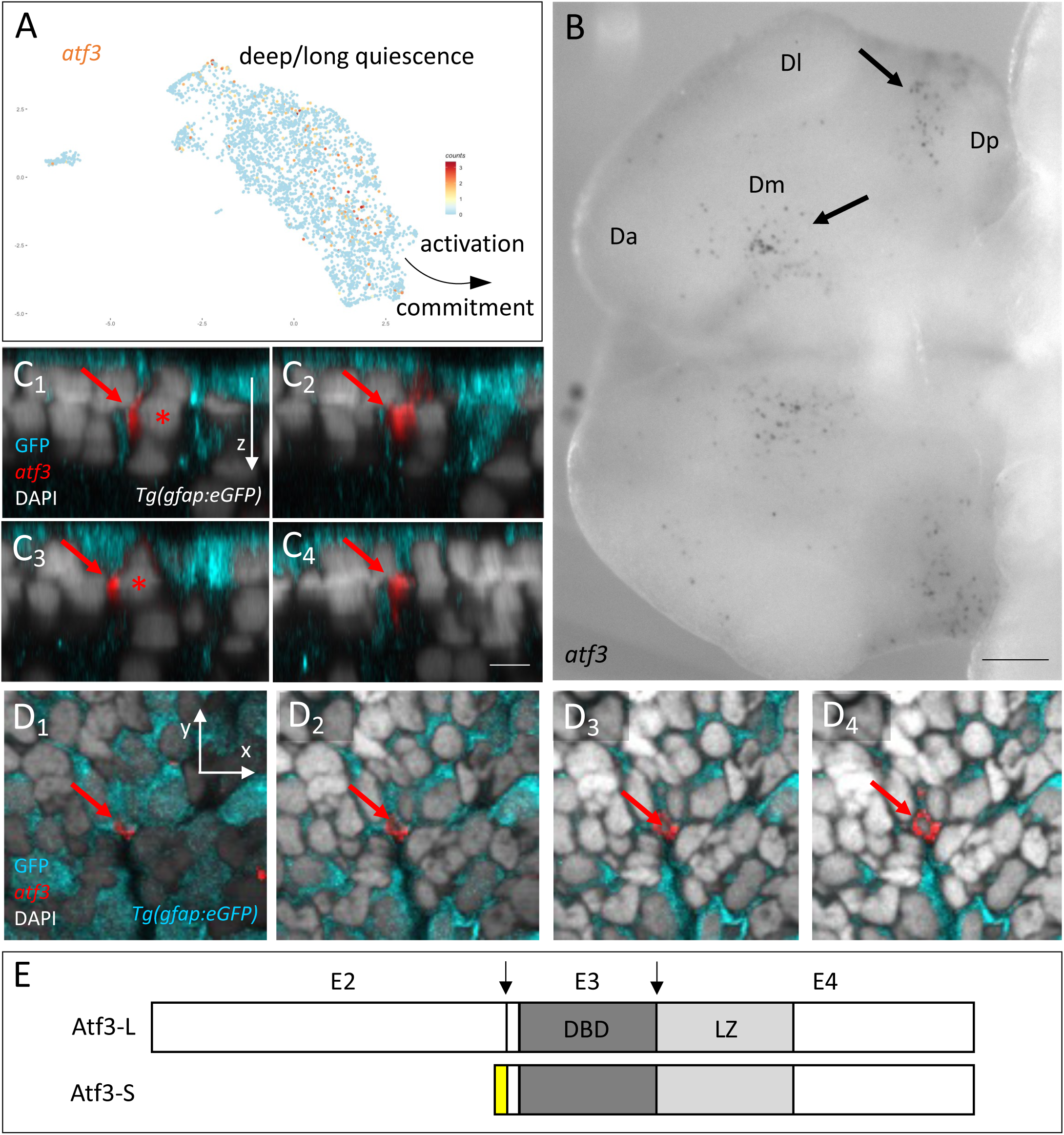
*atf3* is expressed in a subset of NSCs in the adult pallium. **A.** *atf3*-expressing cells (orange dots, UMI values color-coded) positioned on the scRNAseq UMAP of adult quiescent NSCs (gray dots). The positions of NSCs closest to activation and/or neurogenesis commitment and NSCs in a deep/long quiescence phase are indicated (Morizet et al., 2023). **B.** Expression of *atf3* revealed by whole-mount ISH in the adult pallium (dorsal view, anterior left) with the *atf3-L* probe (E). Da, Dl, Dm, Dp: anterior, lateral, median and posterior regions of the pallium, respectively. Arrows to *atf3*-expressing cells in Dm and Dp. **C1-C4.** Optical cross sections of a *Tg(gfap:eGFP)* pallium stained in whole-mount for GFP (cyan), *atf3* transcripts (red) and DAPI (ventricular surface up). The sections are ordered from C1 to C4 along the anteroposterior axis. Red arrows to an *atf3*-positive, GFP-positive cell (red star to the nucleus). **D1- D4.** Optical horizontal sections of the same pallium as in C, ordered from D1 to D4 from superficial to deeper locations. Red arrows to the *atf3*-positive, GFP-positive cell. **E.** Atf3 protein isoforms predicted from adult pallial transcripts (L: long, S: short). The position of coding exons E2-E4 is indicated, as well as splice junctions for the L form (arrows). DBD: DNA-binding domain; LZ: leucine zipper domain. The yellow box indicates the different N-terminus of the S form due to alternative splicing. Scale bars: B: 100µm, C1-4 and D1-4: 5µm.

### Atf3 is necessary for the direct neuronal differentiation of adult pallial NSCs, and impacts physiological *Cas^CRE^Atlas* fate decisions

Next, we combined gain- and loss-of-function experiments in adult pallial NSC in vivo to test whether Atf3 impacts NSC fate. Electroporation of a nlsGFP-encoding construct driven by the ubiquitous *pCMV* promoter upon injection into the cerebral ventricle highlights different fates at short time scale (2 days post-electroporation -dpe-): a large majority of ventricular cells (with a radial morphology and expressing Sox2, likely NSCs), and a minority of delaminating cells (with basally displaced nucleus and ventricular attachment, likely NPs) and of Sox2-negative parenchymal cells (interpreted as neurons) (Fig.5A-E). When electroporated under the same conditions, *pCMV:atf3-S-nlsGFP* had no effect on cell fate (Fig.5E) while *pCMV:atf3-L-nlsGFP* significantly increased parenchymal cells at the expense of ventricular cells (Fig.5D). We conclude that forced expression of Atf3-L leads to the direct delamination of ventricular cells. Because this is reminiscent of non-apoptotic Cas3*-driven fates, we next assessed whether Atf3-L could induce *Cas^CRE^Atlas*. When electroporated into the *Cas^CRE^Atlas* double transgenic background, we observed that *pCMV:atf3-L-nlsGFP* indeed correlated with Hmg2bmCherry expression, although only in a small minority of GFP-positive cells at 7 dpe (Fig.5F, white arrows). Given the low number of electroporated cells, and the low number of ventricular *Cas^CRE^Atlas* cells, we believe that double positive cells are unlikely to result from chance. Their low occurrence is however puzzling. Cas3* may be induced but at levels too low or too transient to generate enough productive Cre recombinase. Alternatively, Atf3 may require a specific context to act upstream of Cas3* in the system studied here. It is to note that the hierarchical position of Atf3 relative to Cas3* has been reported to vary (see Discussion).

**Figure 5.**
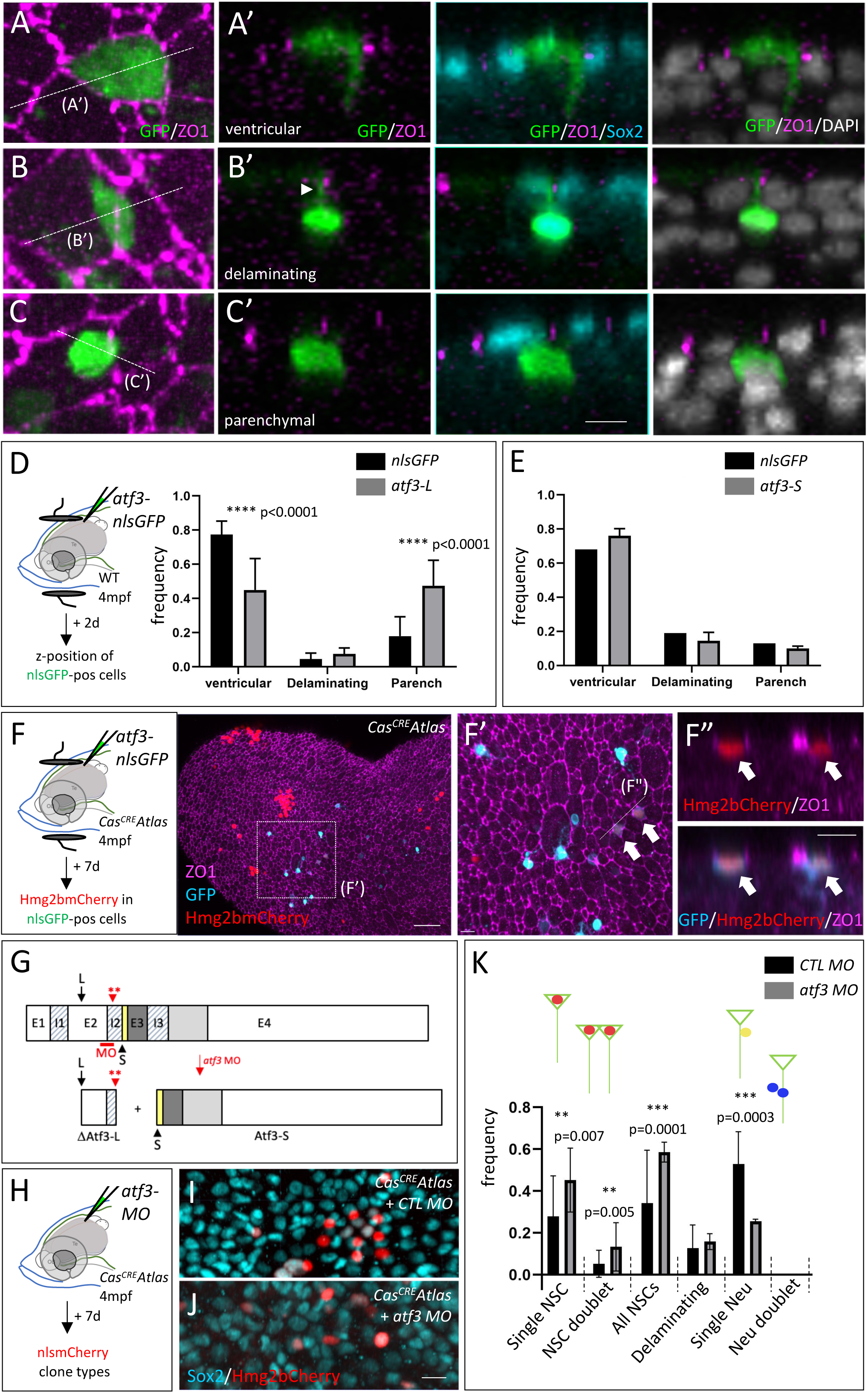
Atf3 can drive NSC delamination and is needed for the expression of the *Cas^CRE^Atlas* fate bias. **A-E.** Cell fates expressed by adult pallial ventricular cells at 2 days-post-electroporation (dpe). **A- C’.** Visual categorization of the three possible fates, illustrated in whole-mount pallia immunostained for GFP (green, electroporated construct), ZO1 (magenta) and the progenitor marker Sox2 (cyan), and counterstained with DAPI (gray). A-C: dorsal (apical) views, A’-C’: cross sections as indicated in A-C. The arrowhead in B’ points to the apical attachment of a delaminating cell. **D,E.** Experimental scheme and quantification of cell fate categories upon electroporation of plasmids for *atf3-L-P2A-nlsGFP* versus *nlsGFP* (D) and *atf3-S-P2A-nlsGFP* versus *nlsGFP* (E). p values: Statistical analysis performed by Chi-square test, 95% confidence D: *nlsGFP* n= 214 cells (4 hemipallia), *atf3-L-P2A-nlsGFP n=69 (3 hemipallia)*, E: *nlsGFP* n= 62 (2 hemipallia), *atf3-S-P2A-nlsGFP* n=121 (3 hemipallia) **F-F”.** Some NSCs overexpressing Atf3-F in *Cas^CRE^Atlas* double transgenic adults are also Hmg2bmCherry-positive after 7 days of chase. Experimental scheme (left) and whole mount pallium processed for triple immunohistochemistry for ZO1 (magenta), GFP (cyan, electroporated construct) and Hmg2bmCherry (red). F’ is a high magnification of F (dorsal views, anterior left) and F” a magnified cross section at the level indicated in F’. Double-positive cells indicated by white arrows. **G.** Exon-intron structure of *atf3* (top) and predicted proteins (bottom) produced upon splice blockade by *atf3-MO* (red bar). Positions of the different exons (E1-E4) and introns (I1-I3) are indicated, as well as the ATG positions for Atf3-L (L, arrow) and Atf3-S (S, arrowhead). The red arrowhead indicates the position of a double stop codon (**) in I2. The color code is the same as in Fig.3C’’’. **H-K.** Blocking Atf3-L function impairs the expression of the *Cas^CRE^Atlas* NSC fate bias. **H.** Experimental scheme. The *atf3-MO* was injected into the forebrain ventricle of *Cas^CRE^Atlas* double transgenic adults and Hmg2bmCherry clone types were quantified after 7 days. **I,J.** Whole-mount *Cas^CRE^Atlas* pallia immunostained for Hmg2bmCherry (red) and Sox2 (cyan) 7 days after injection of the control MO (I) or *atf3-MO* (J) (dorsal views). **K.** Quantification of clone types. “All NSCs” are the sum of “single NSCs” and “NSC doublets”; after this short chase, neurons we only observed as single neurons (the value for “Neu doublets” is null in both conditions). p values: Statistical analysis performed by contingency Chi-square test, 95% confidence, n=6 hemipallia per condition, *CTL* MO: n=82 clones, *atf3* MO: n=43 clones. Scale bars: A-C: 8 µm, F-F’: 50µm, F”: 10 µm, I-J: 8 µm.

We next aimed to test whether Atf3 was required for NSC direct differentiation/delamination. The very low number of *atf3*-expressing NSCs at any time, and our impossibility to track them with currently available tools, did not permit addressing this point in the general context of the entire NSC population. Thus, we more specifically determined whether Atf3 was necessary for direct neuronal differentiation downstream of Cas3*, by testing the effect of blocking Atf3-L function *in vivo* in the *Cas^CRE^Atlas* double transgenic context. We designed a vivo-morpholino (MO) directed against the exon2-intron2 boundary of *atf3*, predicted to generate a truncated Atf3-L protein devoid of its DNA-binding and Leucine zipper domains (Fig.5G). This prediction was validated in embryos (Fig.S2B). This MO should not affect the production of the Atf3-S isoform, the start codon of which is located 3’ to the MO position. The *atf3* vivo-MO, or a control vivo-MO, were injected into the cerebral ventricle of *Cas^CRE^Atlas* double transgenic adults, and *Cas^CRE^Atlas* fates were analyzed after 7 days in whole-mount pallia (Fig.5H-J). Specifically, Hmg2bmCherry-positive clones were categorized when located within the first 1-2 cell rows below the ventricular surface (corresponding to the z-position of NSC-progeny cells generated during 7 days). We found that the *atf3* vivo-MO induced a significant increase in the proportion of clones composed of NSCs only, at the expense of the generation of single neurons (Fig.5K). These results together indicate that Atf3-L expression is necessary for lineage bias, either downstream of or in parallel to physiological non-apoptotic Cas3* events.

### Experimentally induced Cas3/Cas7 activation events drive direct neuronal production from adult NSCs in vivo

The results above indicate that Cas3/Cas7 activation and Atf3-L are, together, linked with the specific NSC fate choice of direct neuronal differentiation under physiological conditions. To further address the relevance of this regulatory process, we first tested whether experimental stimulation of Cas3/Cas7 activity in adult fish could modify NSC fate. CPT was used as an inducer. Incubation in CPT triggered Cas3* induction and was followed by NP death at larval stages (as revealed by Tg(*ubi:secA5-mVenus*) reporter, in which Annexin 5-mVenus expression serves as a marker for cell death) (van Ham et al., 2010) (Fig.S3A-B_1_). In striking contrast, pallial Cas3* cells were seen instead to delaminate when CPT was injected into the cerebral ventricle in adult animals (Fig.6A-C_1_’). To track their fate at longer term, CPT was applied to *Cas^CRE^Atlas* adults and clone types were assessed (Fig.6D-H). We found an increase of clones composed of single neurons (blue in Fig.6F’,H) at the expense of clones made of single NSCs (red in Fig.6F’,H), thus mimicking the fate observed for physiological *Cas^CRE^Atlas* events in the adult pallium.

**Figure 6.**
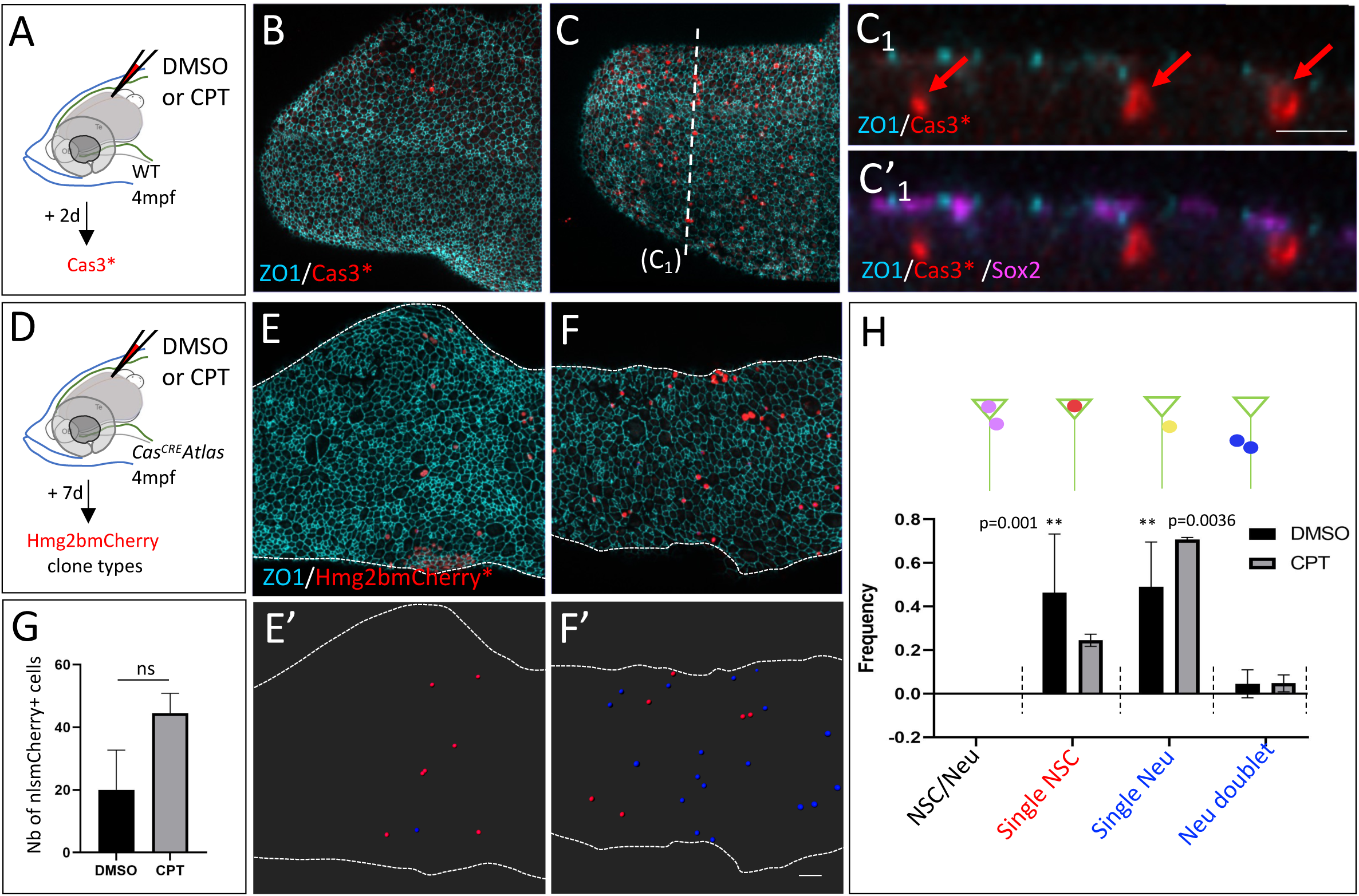
Experimentally induced Cas3* events drive direct neuronal differentiation in the adult pallium. **A-C_1’_.** Induction of Cas3* by CPT (C) compared to DMSO (B) injected into the cerebral ventricle (A), revealed by IHC in adult pallia at 2 days post-injection. **B,C.** Dorsal whole-mount views of the right hemisphere, anterior left, processed in IHC for ZO1 (cyan), Cas3* (red) and Sox2 (magenta, only shown in C’_1_). C_1_ is a cross-section at the level indicated in B, red arrows to delaminating Cas3*-positive cells. **D-G.** Experimental scheme and quantification of clone fate categories in *Cas^CRE^Atlas* adults upon intracerebral injection of CPT (F,F’) compared to DMSO (E,F’) at 7 days post-injection. **D.** Experimental scheme. **E,F.** Dorsal whole-mount views of the right hemisphere, anterior left, processed in IHC for ZO1 (cyan) and Hmg2bmCherry* (red). **E’,F’.** Segmentation of clones generated within 7 days (located at and immediately below the pallial ventricular surface), color-coded (red: NSCs, blue: parenchymal cells, identified as neurons). **G.** Quantification of Hmg2bmCherry-positive cells located within 2-3 cell rows from the ventricle at 7 days post-treatment. Statistical analysis performed by Welsh’s t test: not significant (p=0.17). **H.** Quantification of clone types at 7 days post-treatment. Statistical analysis performed by contingency Chi-square test, 95% confidence. 4 hemipallia each condition. DMSO, n=37 clones, CPT n=89 clones. Scale bars: B,C,E,F,E’,F’,C1,C’1: 30µm.

### Non-apoptotic Cas3 activation events are seldom recruited upon NSC irradiation but contribute to the generation of neurons during lesion repair

Given that Cas3/Cas7 activation events can drive NSC fate change in the adult pallium, we next tested whether such events were recruited under challenged conditions known to impact NSC state or fate. qNSCs have been described as radiation resistant, possibly thanks to an efficient mechanism of DNA repair (Barazzuol et al., 2019; Hellström et al., 2009; Mineyeva et al., 2019). However, radiation-induced differentiation was also described (Konirova et al., 2019; Schneider et al., 2013). We used Xray irradiation of live adults (3mpf) to further support this observation and test whether radiation resistance could also be accompanied by a NSC fate change *in vivo*. Short (1-hour) treatment with a low radiation dose (5 Gy) induced γH2AX-positive foci in NSC/NP nuclei, indicative of double strand DNA breaks and the recruitment of the repair machinery (Fig.7A-B’). This process was transient and completed by 2 hours post-treatment (Fig.S4) with no visible effect on NSC fate and was observed until very high irradiation doses (not shown). At 40 Gy, a low number of cells located very close to the ventricular surface turned positive for Cas3* after a 24-hour chase (Fig.7C-D_1_). These cells displayed a delaminating profile with a cell body partly displaced into the parenchyme (Fig.7D_2_-D_3_) but sometimes keeping a ventricular attachment (Fig.7D_2-2’_) and were Sox2-negative. Although we did not ascertain cell survival at later chase times, these results suggest that, upon irradiation at high dose, a few Cas3* events are induced and correlate with the first steps of neuronal commitment, namely parenchymal relocalization and the loss of expression of the progenitor marker Sox2. The broad range of low to moderate irradiation schemes however lead to repair and do not recruit Cas3*.

**Figure 7.**
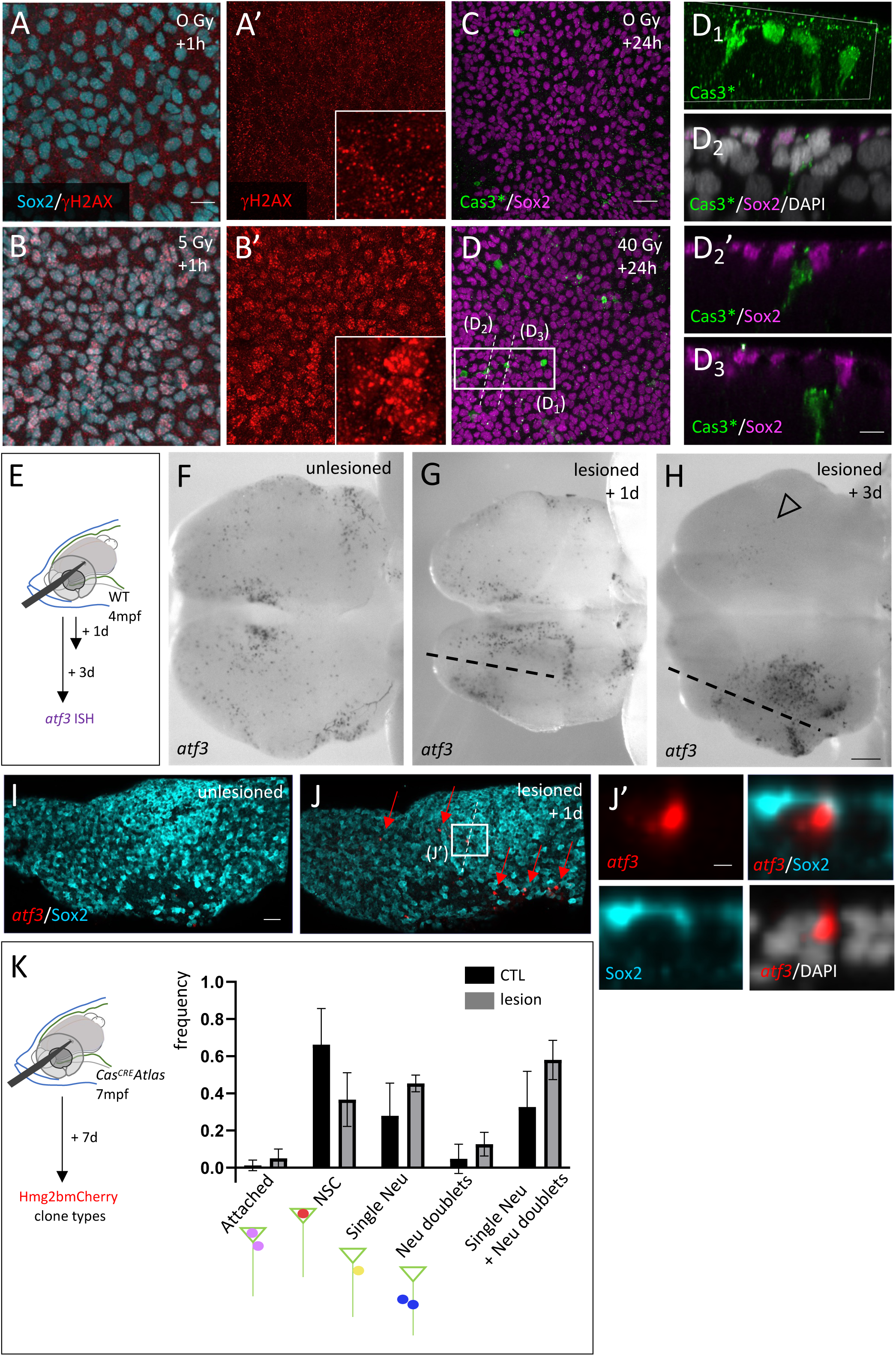
Differential recruitment of *atf3*-positive and Cas3* events under irradiation versus mechanical lesion. **A-D_3_.** Effect of X-rays on NSCs *in vivo*. **A-B’.** γH2AX DNA repair foci (red) in NSCs/NPs (Sox2+, cyan) revealed by whole-mount IHC on adult pallia under control (A,A’) or irradiation conditions (B,B’) (5 Gy + 1hr chase). **C-D_3_.** Casp3* induction (green) in NSCs/NPs (Sox2+, magenta) revealed by whole-mount IHC on adult pallia under control (C) or irradiation conditions (D) (40 Gy + 24hr chase). D_1_ is a high magnification 3D rendering (Imaris) of the area boxed in D, and D_2_-D_3_ are sections across Cas3* cells along the dotted lines in D. The visible channels are color-coded. **E-K.** Induction of *atf3*+/Cas3* events by mechanical lesion. **E-J.** *atf3* expression revealed by chromogenic ISH (F-H: blue, I-J’: red) on whole-mount pallia 1 day (G, I, J) or 3 days (H) following stab-wound injury (E) compared to uninjured brains (F, I). Dorsal views, anterior left, only one hemisphere shown in I, J. Dotted lines: lesion trajectories; open arrowhead: downregulated *atf3* expression in the contralateral hemisphere at 3dpl. J’ is a high magnification section across an *atf3*-positive cell along the dotted lines in J. The visible channels are color-coded. **K.** Experimental scheme (left) and quantification (graph) of the proportion of *Cas^CRE^Atlas* clone types in control versus lesioned hemispheres at 7dpl. Statistical analysis by contingency chi-square test: no significant difference between the proportion of the different *Cas^CRE^Atlas* clone types between lesioned and unlesioned hemispheres. Control 11 hemipallia, n=139 clones, Lesion 3 hemipallia, 43 clones. Scale bars: A,A’, B, B’: 10µm, C-C’: 20µm, D1-D3, D’: 10µm, F-H: 100µm, I-J: 30µm, J’: 10µm.

Mechanical lesions applied to the adult pallium lead to NSC recruitment into division for neuronal repair (Baumgart et al., 2012; Kishimoto et al., 2012; Kroehne et al., 2011; März et al., 2011), accompanied with a partial fate shift towards symmetric neurogenic divisions (Barbosa et al., 2015). Absolute numbers of direct differentiation events were not reported in the latter study. We tested whether *atf3*- positive or Cas3*/Cas7* events could be involved in the response to lesion. We found that *atf3* expression was induced around the lesioned ventricular zone starting at 1 day-post-lesion (dpl) and very prominently at 3dpl (Fig.7E-H, asterisk). It was also massively downregulated in the contralateral hemisphere at 3dpl, a phenomenon not reported yet for other lesion-responsive genes (Fig.7H, open arrowhead). Like under physiological conditions (Fig.4C1-D4), *atf3*-positive cells in lesioned pallia displayed a delaminating morphology (Fig.7I-J’). We next used *Cas^CRE^Atlas* to track the fate of Cas3*/Cas7* events in lesioned context. At 7dpl, *Cas^CRE^Atlas* fates recorded in lesioned hemispheres did not detectably differ, qualitatively and in proportion, from those recorded under physiological conditions (Fig.7K and Fig.5H-K), thereby contributing to reparative neurogenesis via their enhanced direct neuronal production.

## Discussion

In this work, we focus on the adult NSC fate of direct neuronal conversion and bring together several observations in link with this fate. First, we provide a quantitative, morphometric and molecular characterization of delamination/neuronal differentiation events in the NSC/NP population of the adult pallium. This analysis suggests that direct NSC conversions follow a classical delamination process. Second, we demonstrate that non-apoptotic events of Cas3/Cas7 activation physiologically occur in NP/NSCs during development and homeostasis of the zebrafish pallium and, at adult stage, are preferentially associated with the NSC fate choice of direct neuronal differentiation. We identify the transcription factor Atf3 to be necessary downstream of or in parallel to Cas3*/Cas7* for the full expression of this fate. Third, we analyze whether Cas3*/Cas7* events can be recruited to trigger direct neuronal conversion under non-physiological conditions. We show that Cas3*/Cas7* events are induced in adult pallial NSCs in response to mechanical lesion and contribute to the formation of neurons for repair. Together, our results provide a first molecular insight, with lineage and functional tracking in vivo, into the important NSC fate decision of direct neuronal conversion. This process balances fate choices to preserve homeostasis of the pallial NSC population over a lifetime (Than-Trong et al., 2020). We propose that adding such a regulatory level beyond NSC division fate choices could add flexibility, in time and space, to the control of NSC numbers and neuronal production.

The existence of direct neuronal conversion of NSCs, i.e. their acquisition of a neuronal fate in the absence of division, can be postulated from clonal tracing in the adult mouse SGZ (Bonaguidi et al., 2011), and has been observed using intravital imaging in the adult zebrafish pallium (Barbosa et al., 2015; Dray et al., 2015; Than-Trong et al., 2020). The *Tg(gfap:ZO1-mKate);Tg(deltaA:egfp)* intravital imaging dataset allowed us to establish the morphodynamic and molecular characteristics of delaminations and, among those, of those occurring late post-division (at least 14-16 days and up to 41 days in our movies). Our dataset includes NSCs and NPs that cannot be distinguished in intravital imaging but our estimations (Fig.S1) make it highly likely that NSCs are included among the recorded events, and our combined observations suggest that direct NSC conversions follow a classical delamination process. The fact that delaminating NSCs do not undergo apoptosis and convert into neurons, although not directly assessed here, is itself supported by a set of arguments from other works: apoptosis is not observed in the adult pallium under normal conditions (Barbosa et al., 2015; Webb et al., 2009), neurons are the sole parenchymal fate of pallial NSCs (Furlan et al., 2017; März et al., 2010), and delaminating NSCs can be traced to express neuronal markers (Barbosa et al., 2015).

Our analysis further suggests that *Cas^CRE^Atlas* captures a measurable fraction of these events, which are characterized by the expression of a specific molecular pathway involving Cas3*/Cas7* activation and Atf3. Several arguments support this conclusion. First, the intravital imaging dataset suggests a number of direct conversion events of the same order of magnitude as the number of *Cas^CRE^Atlas* clones quantified over the same period (NSC delaminations in intravital imaging: likely a few dozens of events in ∼800 NSCs in 40 days; *Cas^Cre^Atlas*: 37-46 events in ∼1000 NSCs in 56 days). Second, *Cas^CRE^Atlas* activation is driven in NSCs and generates persisting clones (Fig.3). Third, CPT treatment reveals that induction of the apoptosis cascade is sufficient to trigger NSC delamination (Fig.6), while Atf3 blockade impairs the expression of the direct neuronal conversion fate (Fig.5G-K), together suggesting that a cascade involving Cas3* and/or Cas7* and Atf3, more than being a simple marker, actually drives the neuronal conversion fate. Obviously, a missing step that would connect our morphometric description with the Cas3*/Cas7*/Atf3 cascade would be the direct tracking of the latter molecular events in an intravital imaging approach. This is however not possible with our current tools, notably because Hmg2B-mCherry needs several days to be directly visible by fluorescence, which would bypass the initial steps of the fate process. An NSC transcriptomic state possibly linked with direct conversions was recently proposed based on in silico analyses of single-cell RNAseq data (Mitic et al., 2024). This interpretation remains to be validated with lineage and functional assays in vivo, but its relationship with Cas3*/Cas7* and Atf3 could be interested to assess.

Beyond this restriction, an important finding of our work is, in itself, the fact that non-apoptotic Cas3/Cas7 activation events are physiological components of NSC population fates: such events do occur in NPs of the developing pallium and in NSCs of the adult pallium, and bias NSC fates towards the direct generation of neurons (Fig.3). To our knowledge, this is the first demonstration of non-apoptotic Cas3/Cas7 activation events being associated with a specific SC fate choice in vivo. In this context, it is important to keep in mind that our work does not imply that this NSC fate is the consequence of a “rescue” of NSCs from death, but simply that this fate is associated with Cas3/Cas7 activation. At the molecular level, the *Cas^CRE^Atlas* principle and the functional experiments of the present work provide some information on the effectors or facilitators of this process. The cleavage site of the *Cas^CRE^Atlas* construct is also recognized by Cas7* and we cannot exclude a contribution of this effector Caspase to the *Cas^CRE^Atlas* signal. *casp7* is not expressed in the larval brain (Spead et al., 2018), arguing for a Cas3*-specific cleavage at larval stages (Fig.2). However, transcription of both *casp3* and *casp7* is also detected in our scRNAseq dataset of quiescent NSC (Fig.S2A). Nevertheless, we observe a correlation between Cas3* IHC and *Cas^CRE^Atlas* events in the number and spatial distribution in response to CPT (Fig.6), strongly suggesting that *Cas^CRE^Atlas* reads, at least in part, Cas3*. Finally, when the production of the transcription factor Atf3 is abrogated, *Cas^CRE^Atlas* is still induced in NSCs, but its associated fate bias is abolished (Fig.4K), suggesting that Atf3 is either a mediator of non-apoptotic Cas3*/Cas7* events or a parallel and converging actor. There remain however several missing steps in our molecular understanding of this NSC fate. The relationship between non-apoptotic Cas3*/Cas7* and Atf3 is not a simple hierarchy, as overexpressed Atf3 is also capable to some extent of inducing *Cas^CRE^Atlas* (Fig.5F). This is reminiscent of previous observations, where Atf3 was described both upstream and downstream of Cas3*, with highly context-dependent mechanistic links (Lu et al., 2007; Syed et al., 2005). Atf3 targets in adult NSCs remain to be identified, as well as Cas3*/Cas7* effectors, given that blocking Atf3 does not fully abolish direct neuronal differentiation (Fig.5F). Additionally, our work does not identify the functionally relevant Casp3*/Casp7* substrate the cleavage of which will bias NSC fate. We note that intense DNA damage, which can be caused by Caspase activation (upon cleavage of the Caspase-activated DNAse inhibitor) and trigger cell differentiation e.g. in muscle (Larsen et al., 2010), is here efficiently repaired in adult pallial NSCs (Fig.7), suggesting that other Caspase substrates are engaged in NSC fate. Along these lines, it would be interesting to search for NSC fate determinants containing Casp3*/Casp7* cleavage sites. Finally, and importantly, we note that *Cas^CRE^Atlas* is induced in NSCs following other endogenous fates, such as NSC-maintaining divisions (Fig.3D). Thus, Cas3*/Cas7* alone is not sufficient to encode the direct differentiation fate in NSCs and a further level of regulation must exist.

Our results also demonstrate that Cas3*/Cas7* events, and the associated fate of direct neuronal differentiation, can be engaged under non-physiological conditions of NSC stress (e.g. very high doses ionizing radiations) or NSC recruitment for neuronal repair (e.g. upon mechanical lesion). In the latter situation, direct neuronal conversion appears recruited without bias among other neurogenic fates (Fig.7K), and at present we did not identify a situation leading us towards the identification of Cas3*/7*/Atf3 induction pathways. Outside the brain, direct SC differentiation has been reported in adult mouse muscle satellite cells, at low levels under physiological conditions and much increased upon abrogation of Notch signaling or under conditions of regeneration, which disrupt MuSC quiescence (Bjornson et al., 2012; Ismaeel et al., 2023; Mourikis et al., 2012). Notch3 signaling is also a major gatekeeper of NSC quiescence in the adult pallium (Alunni et al., 2013; Chapouton et al., 2010) and whether abrogating this pathway and/or quiescence will result in mobilizing direct neuronal differentiation fates in addition to NSC activation for proliferation remains to be tested. It is to note that neuronal differentiation is not the “default” fate of perturbed NSCs, as, for example, adult zebrafish pallial NSCs will take a fate of differentiated astroglia (and not neurons) when Notch-induced stemness is abrogated (Than-Trong et al., 2018). Along the same lines, irradiation can drive the differentiation of proliferating neural progenitors towards the neuronal (Konirova et al., 2019) but also the astrocytic (Schneider et al., 2013) fates.

The association of caspase events with cell fate changes in a number of physiological contexts (Burgon and Megeney, 2018; Fujita et al., 2008; Janzen et al., 2008), or with cellular remodeling processes such as axonal degeneration or regeneration, dendritic pruning, axonal pathfinding, synaptic plasticity notably in the nervous system (Unsain and Barker, 2015), led to postulate *bona fide* non-apoptotic functions of the caspase pathway. Atf3 itself, in addition to a stress induced factor, is also seen as an adaptive response element driving cellular remodeling under stimulation by a number of signaling pathways (Rohini et al., 2018). Our work adds NSCs to the list of cells whose fate choices can be influenced by non-apoptotic caspase events. Neuronal differentiation and (neuro)epithelial delamination, such as undergone by delaminating NSCs in the adult pallium, both involve major cellular remodeling, such as massive changes in cytoskeletal architecture and intracellular trafficking, nuclear repositioning, constriction of the apical and ciliary membranes, junction disassembly and abscission (Kasioulis and Storey, 2018; Kuijpers and Hoogenraad, 2011). Some of these are shared with the major cellular rearrangements that initiate apoptosis, which may go together with sharing core control pathways. It will of course also be interesting to determine whether the neurons issued from direct neuronal conversion of NSCs have specific structural features and identity.

## Materials and methods

### Fish lines

Wild-type (AB) and *Tg(GFAP:eGFP)* (Bernardos and Raymond, 2006), *Tg(ubi:SecA5)* (van Ham et al., 2010), *Tg(her4:ERT2CreERT2)* (Boniface et al., 2009), *Cas^Cre^Atlas* (see below), *Tg(βactin:lox-stop-lox-hmg2B- mcherry)* (Wang et al., 2011) transgenic zebrafish were used. Embryos/larvae up to 5 dpf were maintained and staged as described (Kimmel et al., 1995). Adult zebrafish were maintained using standard fish-keeping protocols and in accordance with our Institute’s Guidelines for Animal Welfare.

### Plasmids/vectors construction and transgenesis

The *mCD8-DEVD-V5-Cre* (*CDVC*) construct was PCR-generated by fusing in frame the *mCD8-Diap1* region of the plasmid encoding Drosophila *Casexpress* DQVD (Ding et al., 2016) to a *V5-CRE* recombinase cassette and subcloned into the Tol2-kit vector *pME-MCS* to generate *pME-CDVC*. *pME- CDVC* was used to generate the transgenesis vector *pTol2-HCDVC* (*her4:mCD8-diap1-V5-CRE-SV40pA*) using the L/R recombinase reaction and the Tol2 vectors p302, p395, and the 5’ vector *N11*. In the final product, the DEVD caspase site was mutagenized to GSGC to generate the control plasmid *pTol2- HCDVC\** by the Round-the-horn mutagenesis method (https://openwetware.org/wiki/’Round-the-horn_site-directed_mutagenesis). Transgenic lines were made by injecting 1-cell embryos with a mix containing 60 ng/μl of plasmid and 60 ng/μl of *transposase* capped RNA.

The *atf3* cDNAs was amplified from reverse transcribed 16hpf embryo RNA and subcloned into *pSCA*. The *atf3* RNA probe was generated from the long version of the RNA (full coding sequence). The *atf3- P2A-GFP* constructs were generated with the Gibson method and subcloned into the *pCMV5* vector using the NEBuilder^®^ HiFi DNA Assembly Cloning Kit.

### RT-PCR for the validation of efficiency of *atf3* vivoMO

cDNA, extracted from 24hpf embryos, was amplified by RT-PCR using primers *atf3_FL_fwd* and *atf3_FL_rev* and the following cycles: 98°C, 1min; 98°C, 30 sec; the 35 cycles with 98°C, 10 sec; 63°C, 30sec; 72°C, 20Sec; then 72°C, 2 min..

### Immunohistochemistry (IHC)

Brains were dissected in 1X PBS at 4°C, their tela choroida was manually removed and the brains were directly transferred to a 4% paraformaldehyde solution in PBS for fixation. They were fixed overnight at 4°C under permanent agitation. After four washing steps in PBS, brains were dehydrated through 5 to 10 minutes series of 25%, 50% and 75% methanol diluted in 0.1% tween-20 (Merck) PBS solution and kept in 100% methanol (Merck) at −20°C. Rehydration was performed using the same solutions, and then brains were processed for whole-mount immunohistochemistry (IHC). After rehydration, the telencephala were dissected out and subjected to an antigen retrieval step using Histo-VT One (Nacalai Tesque) for 1 hour at 65°C. Brains were rinsed three times for at least ten minutes in a 0.1% DMSO and 0.1% Triton X-100 (Merck) PBS 1X solution (PBT) and then blocked with 4% normal goat serum in PBT (blocking buffer) 4 hours at RT on an agitator. The blocking buffer was later replaced by the primary antibody solution (diluted in blocking buffer), and the brains were kept overnight at 4°C on a rocking platform. The next day, brains were rinsed five to ten times over 24 hours at room temperature with PBT and incubated in a solution of secondary antibodies diluted in PBT overnight, in the dark, and at 4°C on a rocking platform. In some instances, to be able to use the ZO1 dye-coupled antibody in the presence of another primary mouse mAB, secondary antibody free sites were blocked by incubation with 2% mouse serum in PBT for 1 hr before applying the dye-coupled ZO1 antibody. After three rinses in PBT over 4 hours, brains were transferred into PBS. Dissected telencephala were mounted in PBS on slides using 0.5 mm-thick holders. The slides were sealed using a glue gun.

Primary antibodies were used at a final concentration of 1:1000 for chicken anti-GFP and chicken anti BrdU, 1:500 Dye-coupled-ZO1, 1:300 Casp3a, 1:250 for DsRed, 1:200 for Sox2, ZO1, mAb anti GFP, 1:100 for γH2AX. Secondary antibodies were all used at a final concentration of 1:1000.

In situ hybridization was performed as described previously (Bosco et al., 2013; Chapouton et al., 2010; Ninkovic et al., 2005) except for the additional presence of 5% dextran-sulfate during the hybridization phase. For combined ISH and IHC, the ISH was developed using Fast-red. See Table S1 for detailed antibodies and probes used in this study.

### Whole-mount in situ hybridization (ISH) and immunohistochemistry (UHC)

*In situ hybridization* was performed as described previously (Bosco et al., 2013; Chapouton et al., 2010; Ninkovic et al., 2005) except for the additional presence of 5% dextran-sulfate during the hybridization phase. For combined ISH and IHC, the ISH was developed using Fast-red (Sigma, F4648). See Table S1 for detailed probes used in this study.

### Ventricular Micro-injections and Electroporation

Micro-injections into the adult brain ventricle were performed on anaesthetized fish as described (Rothenaigner et al., 2011) except that DNA was injected at the midbrain midline to avoid damaging the pallium. vivoMOs (Gene-tools) were injected at a concentration of 0.125 mM. Micro-injections into 4dpf larval hindbrain ventricle were performed on anesthetized larvae immobilized in 4% methyl-cellulose. For electroporation, plasmid DNA was diluted to 1μg/μl in 0.1 x PBS and injected into the ventricle. Electrodes (TWEEZERTRODES, 5mm Platinum) were placed on each side of the fish head. Fish were then administered two electric pulses (70 V, 50 ms width, 1,000 ms space).

### Drug treatments

4-Hydroxytamoxifen (4-OHT) Treatments and Bromodeoxyuridine Incorporation: 4-OHT (T176, Sigma) treatment was performed as previously described (Mosimann et al., 2011) on *her4:Ert2CreErt2*, *ßactin:LoxSTOPloxhmgbmCherry*. Clonal recombination conditions at 1 mpf were 10 min with 0,5µM 4- OHT as in Than-Trong et al, 2020. They were followed by a 4h pulse of 1mM BrdU. *Cas^Cre^Atlas* were only treated with 1mM BrdU for 4h. Fish were then washed four times, transferred into fresh fish water, and grown as usual until 2-month-old.

Camptothecin treatment: CPT was dissolved as a 10mM stock in DMSO, aliquoted and stored frozen until use. Just before use, an intermediate solution was prepared in DMSO and further diluted 50 times in PBS (for injections) or fish water (for incubations) to reach the working concentration.

### X-ray irradiation

3-5 month old fish where placed into 2-liter cages inside an XRay irradiator (Gulmay CP160/10, 250kV, 12mA) and irradiated for the length of time needed to reach the expected dose (5 Gy= 212 sec). Controls were placed for the same duration in the chamber but left untreated. Following irradiation, fish were kept in 6-liter cages in a 28°C incubator for the desired length of time. Fish were fed once a day and water changed daily if needed. Fish brains were then dissected, their tela choroida was removed, and the brains were then fixed for ISH or IHC.

### Imaging and image analysis

Images of whole-mount immunostained telencephali were acquired on a confocal microscope (LSM700 and LSM700, Zeiss) using a 20X objective or a 40X oil objective (Plan-Apochromat 40x/1.3 Oil M27) and tile images of 4 to 8 z stacks were stitched with the ZEN2009 software. 3D renderings were generated using the Imaris® software (versions 8 and 9, Bitplane). Vertical plane images were extracted when needed. *Cas^CRE^Atlas c*lones were resolved manually using 3D rendering and the slice mode and highlighted in different colors. For figures 7A-D and Supp 3, a nuclear mask was created in IMARIS using the sox2 nuclear staining in order to quantify nuclear H2AX staining. For figure 4D, single planes at different depth were extracted.

For dorsal whole-mount views of the telencephalon (ISH in blue), images were taken using a Nikon macrozoom.

### Statistics

All experimental data were analyzed using Prism software and are expressed as mean ± 95% confidence interval (95% CI). Significance was set at p < 0.05. Comparison of proportions between experimental and control conditions (Figs. 3D, 5D-E, 6H, 7K) were performed via a contingency test based on a Chi2 analysis. For all other comparisons, a Mann-Whitney test (non-parametric test, non-gaussian distribution) was used.

## Acknowledgments

We thank the ZEN team for input, Isabelle Foucher for expert assistance in particular with the generation of *her4:ERT2CreERT2*-driven clones, Emmanuel Than-Trong for his specific re-analysis of 4- OHT-induced clones after a 4-week chase (for comparison with Fig.3D), Laure Mancini for initially producing the movies that were here re-analyzed in Figures 1 and S1 for direct differentiation events, the Institut Pasteur Irradiator service platform (Philippe Casanova and Claire Mallet) for expert assistance with X-ray use, and Emeline Perthame from the Institut Pasteur Bioinformatics platform for her help in the choice of tools for statistical analyses. We thank Denise Montell for sharing constructs, Tjakko van Ham for sharing the *Tg(ubi:SecA5)* line. We are also greatly indebted to Romain Levayer and his lab, Miria Ricchetti and Shahragim Tajbakhsh for insightful discussions and suggestions, and to Romain Levayer for his critical reading of the manuscript.

## Author contributions

Conceptualization: FR, LBC; Methodology: FR, ND, LBC; Funding acquisition: LBC; Project administration: LBC; Supervision: LBC; Writing – original draft: LBC; Writing – review & editing: FR, ND, LBC.

## Data availability

All data used in the analysis will be made available upon request.

---

**Figure S1.**
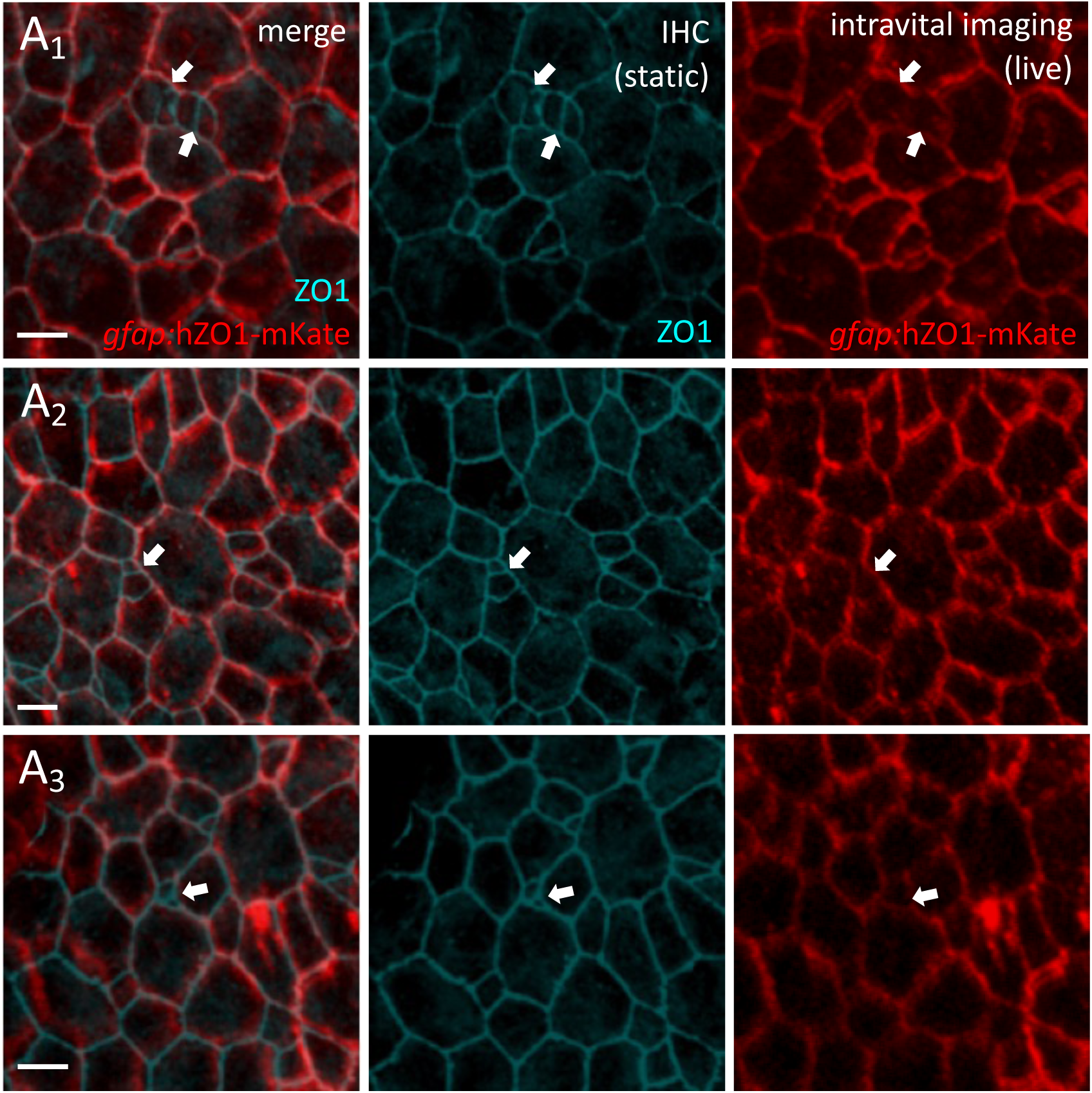
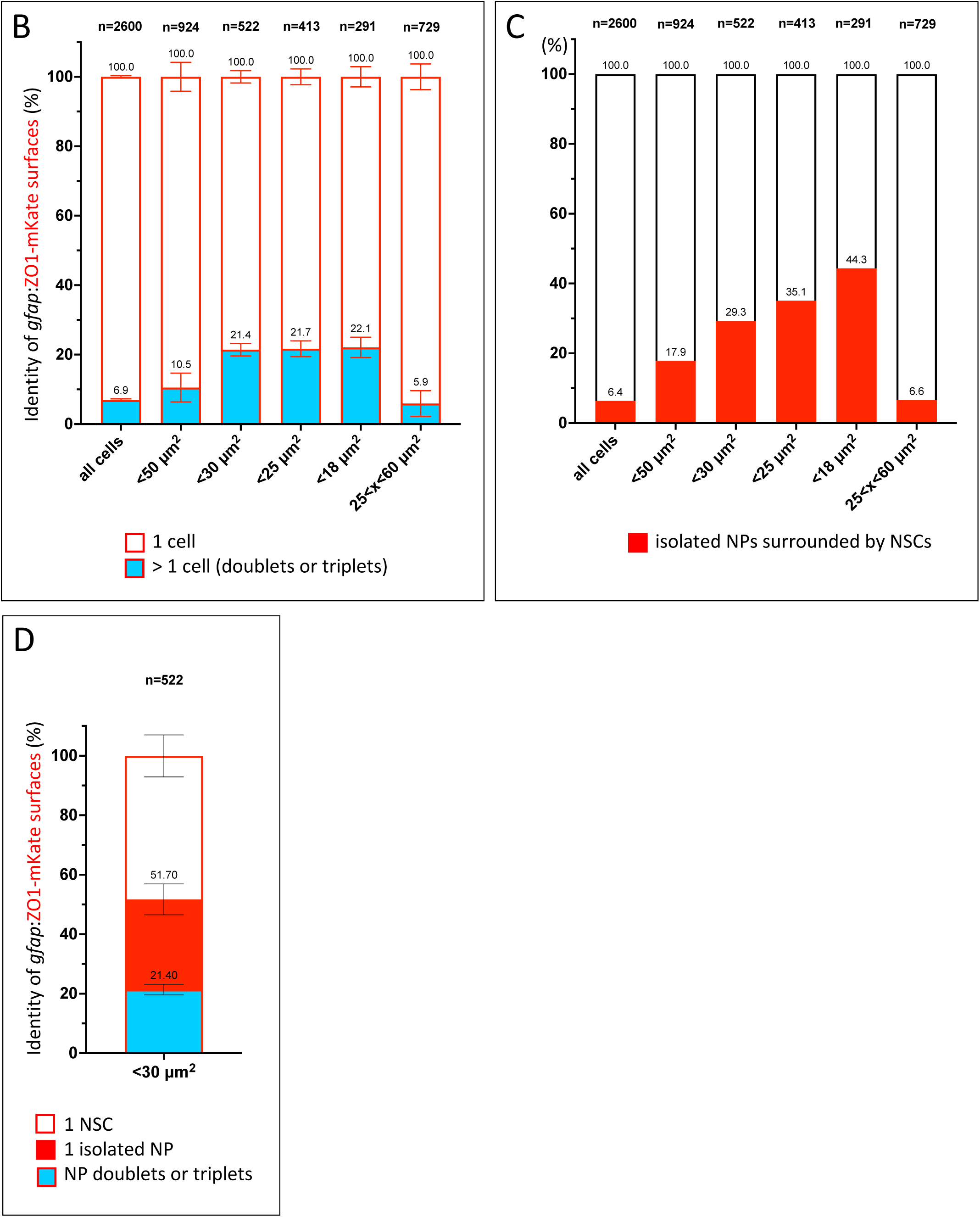
Estimation of the proportion of ZO1-mKate apical surfaces corresponding to NPs in the intravital imaging approach. **A_1_-A_3_.** Three examples of whole-mount overlays between ZO1-mKate fluorescence at a final imaging tp (red) and ZO1 immunohistochemistry (IHC) on the same brains following fixation (cyan) (Da pallial domain). The merge panel shows the unambiguous alignment of the fixed and live images, except for some adjacent NPs (white arrows), which cannot be recognized as distinct apical domains in the live image because expression of the ZO1-mKate fusion protein is driven by the NSC *gfap* regulatory elements. Scale bars: 10 μm. **B.** Proportion of ZO1-mKate surfaces corresponding to NP clusters (cyan) versus single cells (white), estimated when comparing ZO1-mKate (live) and ZO1 (fixed). The percentage is estimated for different cut-offs of apical surface areas (in μm^2^). Da pallial domain, n=2 brains. **C.** Proportion of ZO1 surfaces corresponding to NPs isolated among NSCs, quantified on Tg(*gfap:dTomato*) pallia following ZO1 IHC. The percentage is estimated for different cut-offs of apical surface areas (in μm^2^). Dm pallial domain, n=2 brains. **D.** Compiled from B and C, identity of ZO1-mKate surfaces < 30 μm^2^. 48.3% are NSCs (white), 51.7% are NPs (isolated NPs - red- or NP groups -cyan-).

**Figure S2.**
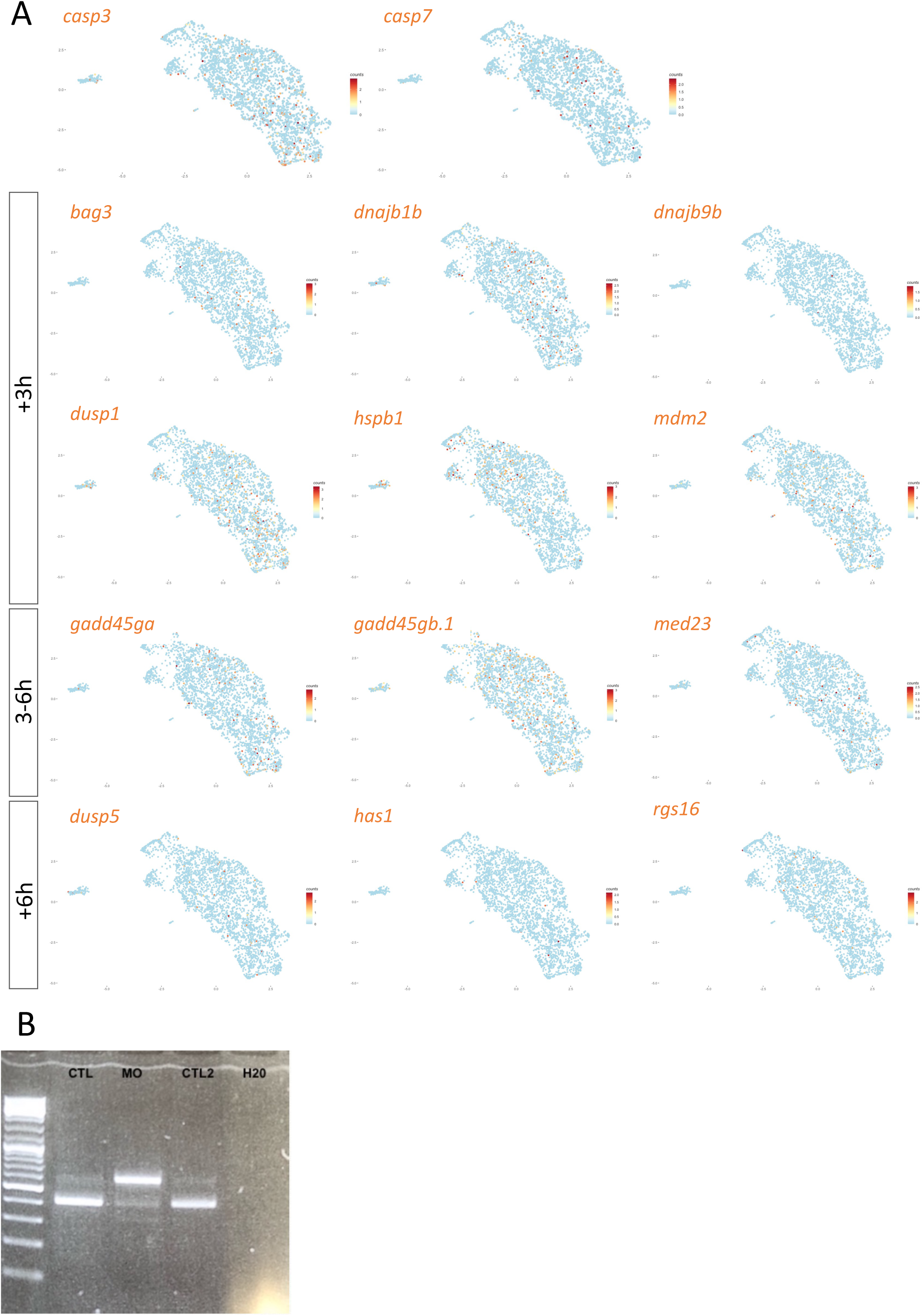
Expression of apoptosis/anastasis mediators in zebrafish adult pallial NSCs and validation of the *atf3* MO. **A.** Cells expressing the genes indicated (orange dots, UMI values color-coded) positioned on the scRNAseq UMAP of adult quiescent NSCs (gray dots). The candidate genes are taken from (Sun et al., 2017; Tang et al., 2017). The positions of NSCs closest to activation and/or neurogenesis commitment and NSCs in a deep/long quiescence phase are as in Fig.2A. **B.** RT-PCR for *atf3* in 24hpf embryos treated as follows: CTL: non-injected, MO: injected at the one-cell stage with 125mM of *atf3* vivoMO, CTL2: injected at the one-cell stage with 125mM of control vivoMO, H20: no RNA.

**Figure S3.**
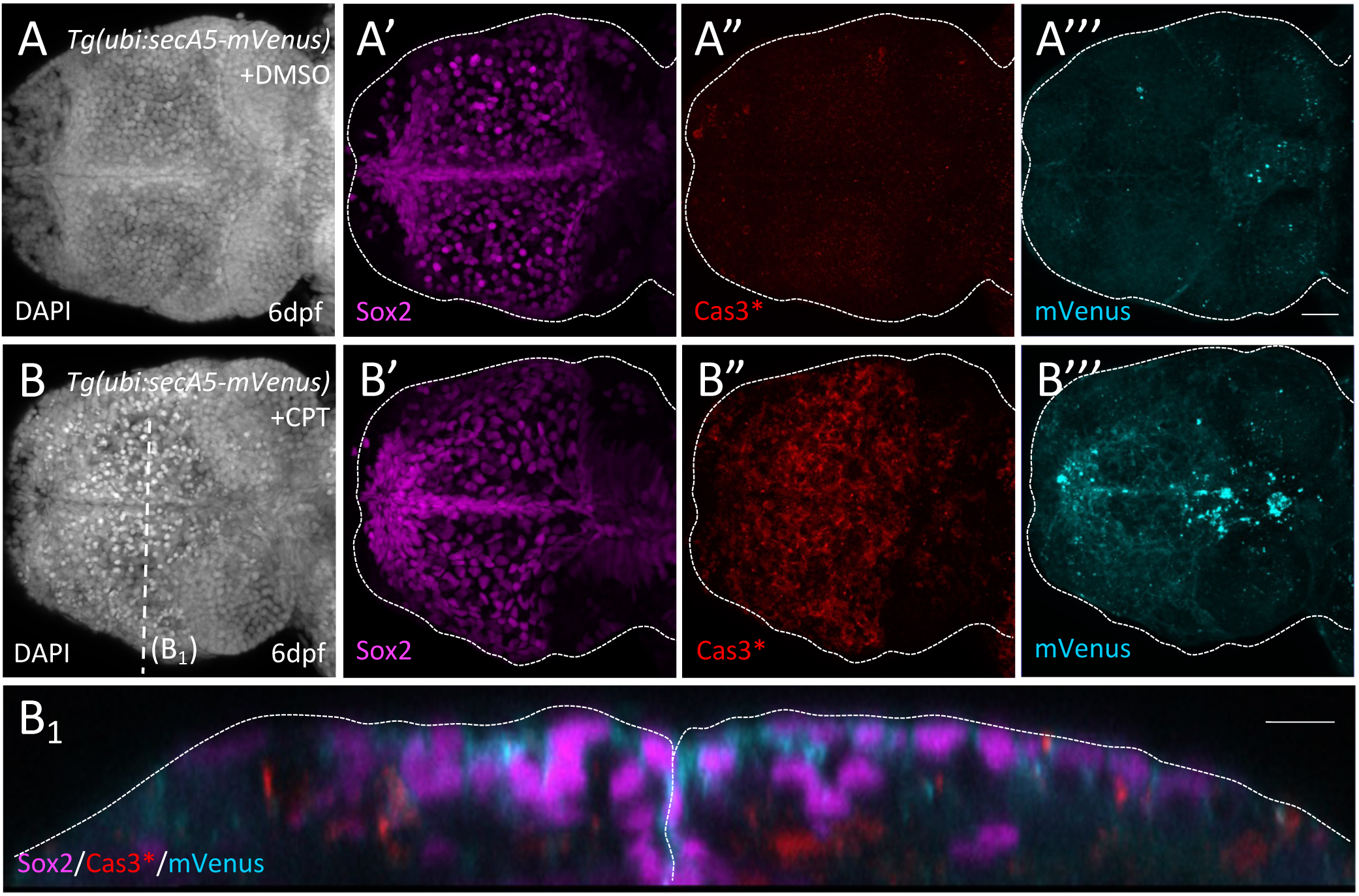
Experimentally induced Cas3* events drive NSC death in the larval pallium. Tg(*ubi:secA5- mVenus*) 6dpf larvae were incubated overnight in DMSO (**A**) or CPT (**B**) (400nM). The brains were dissected and subjected to whole-mount IHC for Sox2 (magenta), Cas3* (red) and mVenus (cyan). B_1_ is an optical cross-section of B at the level indicated. Note that the mVenus staining is ventricular and affects Sox-positive NSC/NPs. Scale bars: A-A3’’’,B-B’’’: 15µm, B1: 10µm

**Figure S4.**
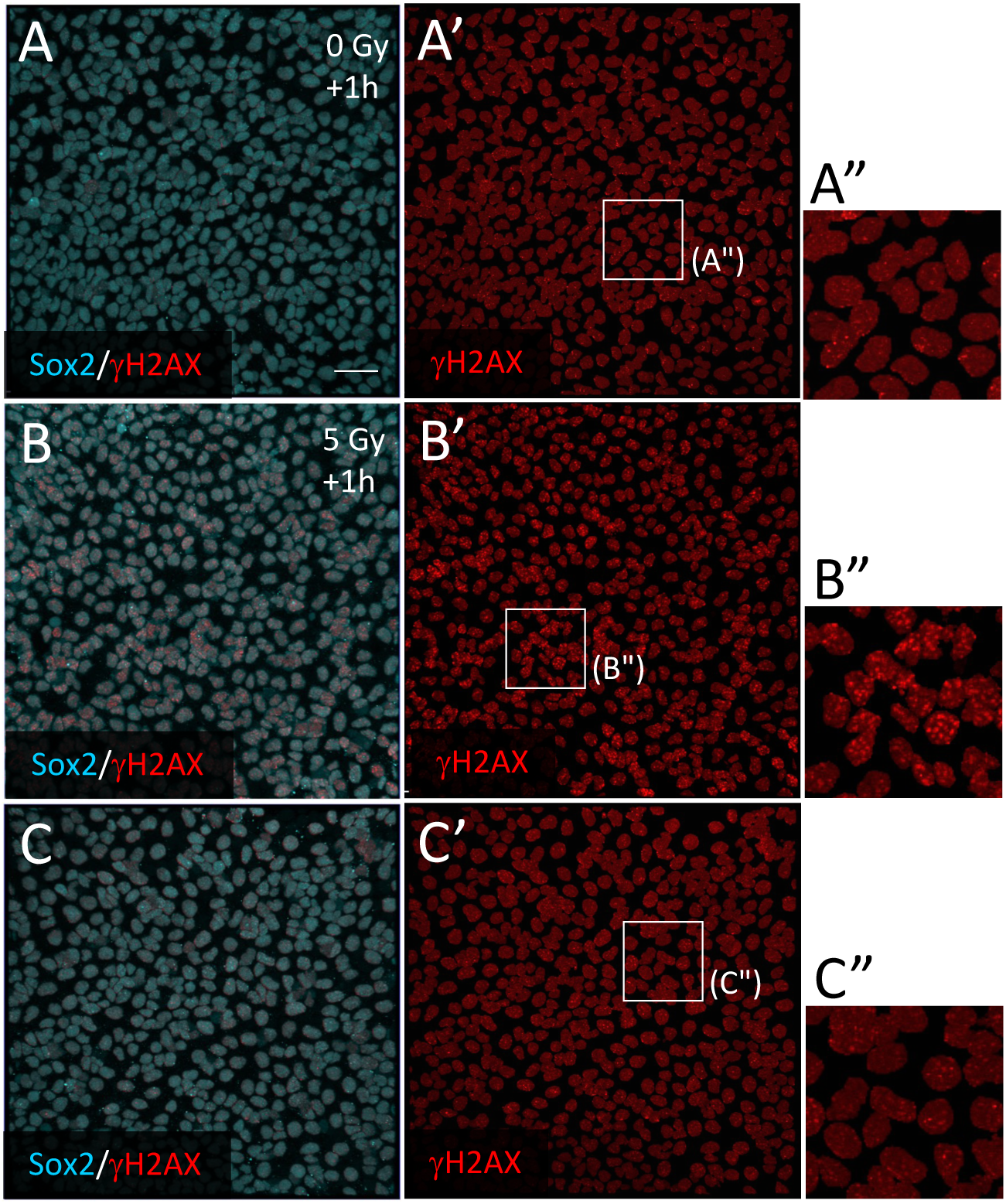
Time course of the effect of X-rays on NSCs *in vivo*. γH2AX DNA repair foci (red) in NSCs/NPs (Sox2+, cyan) revealed by whole-mount IHC on adult pallia under control (A,A’) or irradiation conditions (B-C’) (5 Gy + 1 or 2hr chase). γH2AX DNA repair foci are prominent 1h after treatment but resolved after 2h. Scale bar: 10 µm.

**Table S1:**
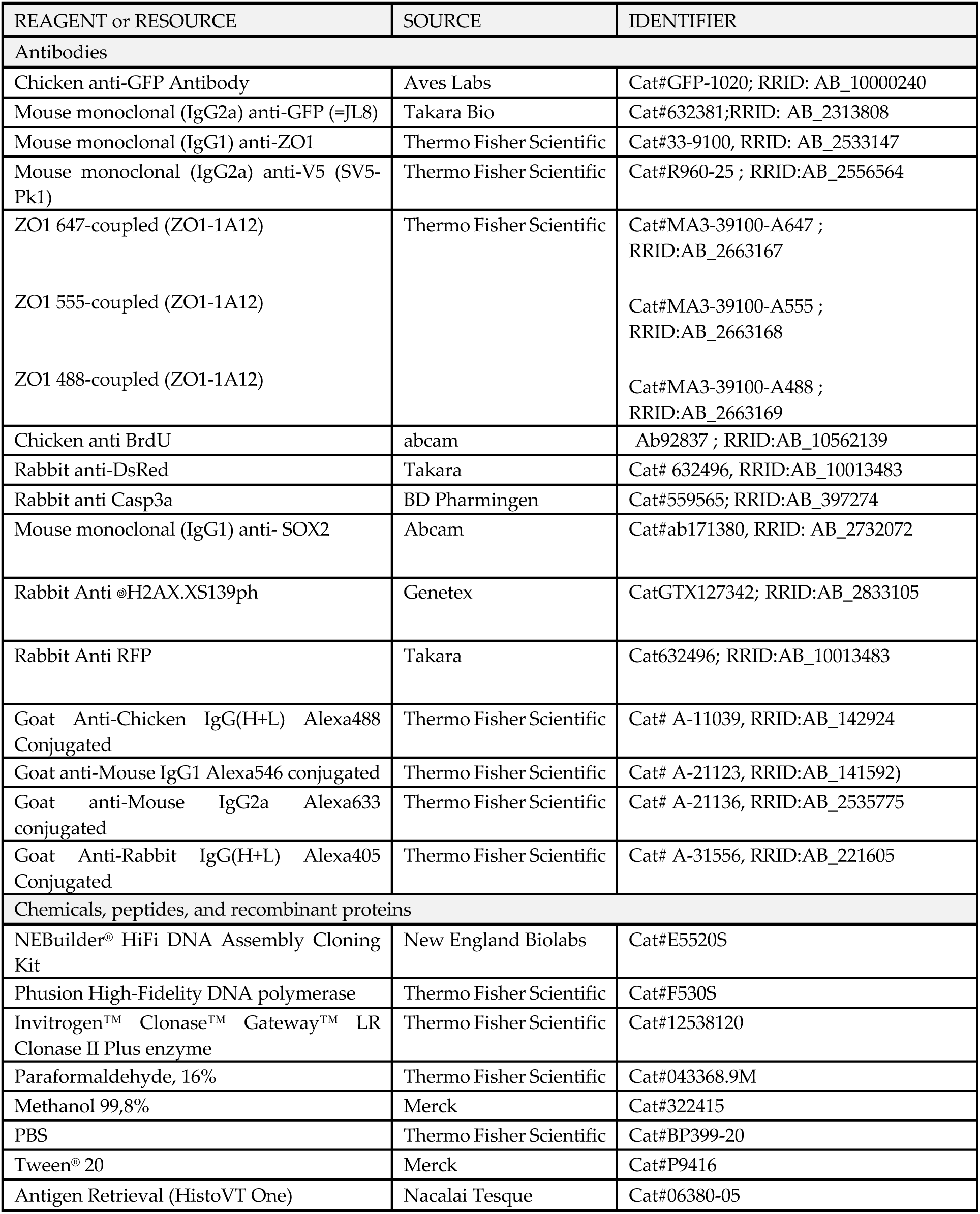
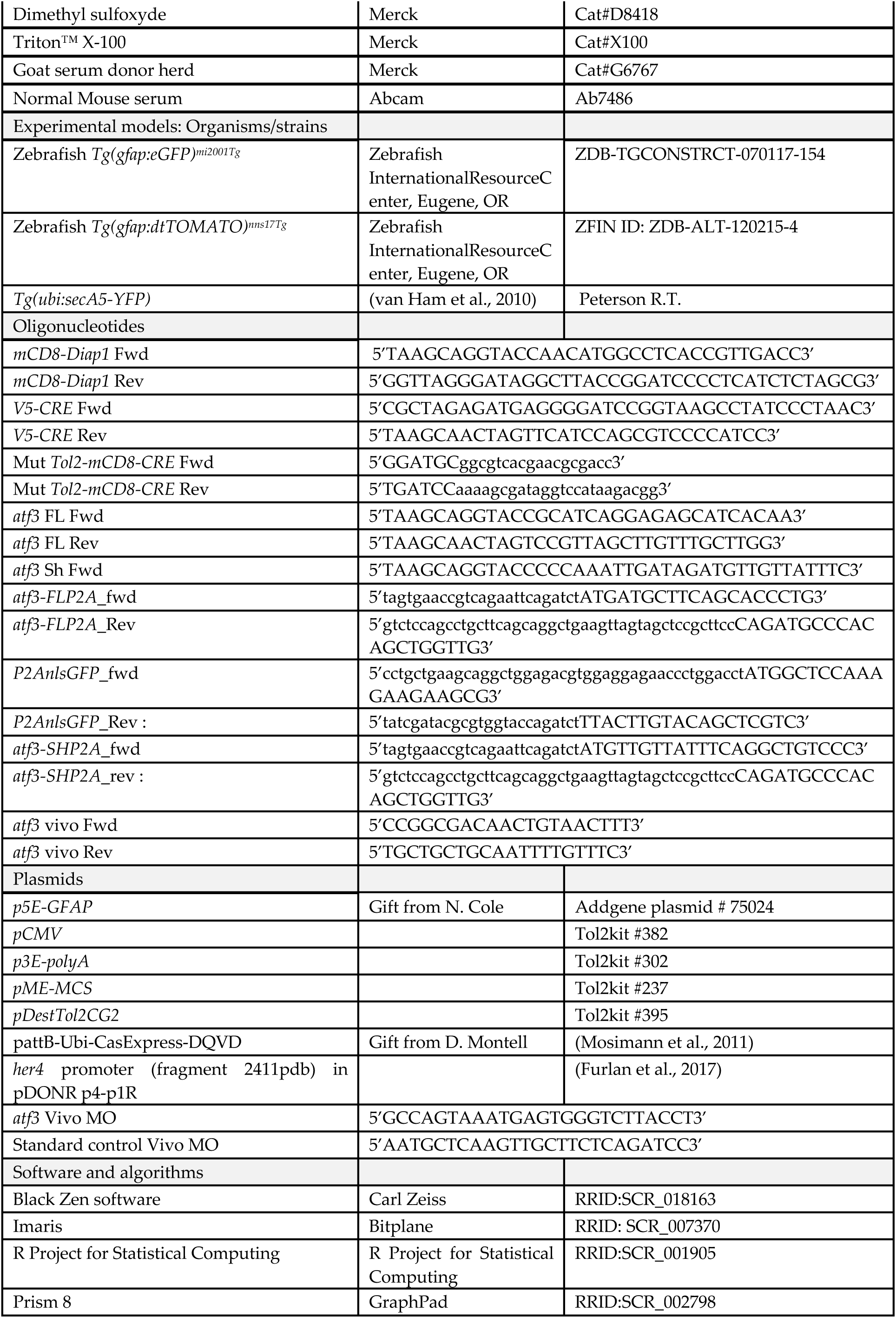
Tools and reagents.

